# Nulliparity affects the expression of a limited number of genes and pathways in Day 8 equine embryos

**DOI:** 10.1101/2022.01.19.476782

**Authors:** E. Derisoud, L. Jouneau, C. Archilla, Y. Jaszczyszyn, R. Legendre, N. Daniel, N. Peynot, M. Dahirel, J. Auclair-Ronzaud, V. Duranthon, P. Chavatte-Palmer

## Abstract

Nulliparous mares produce lighter and smaller foals compared to mares having previously foaled, with effects observed at least until 4 months of age. The need for a first gestation priming for the uterus to reach its full capacity has been proposed to explain this observation. Embryo developmental defects could be hypothesized but effects of maternal parity on the embryo have only been described once, in old mares, thus combining effects of parity and old age. The aim of this study was to determine effects of mare parity on embryo gene expression. Day-8 post ovulation blastocysts were collected from young (5/6 years old) nulliparous (YN, N=6) or multiparous (YM, N=4) non-nursing Saddlebred mares, inseminated with the semen of one stallion. Pure (TE_part) or inner-cell-mass-enriched (ICMandTE) trophoblast were obtained by embryo bisection for RNA sequencing (paired end, non-oriented, Illumina, NextSeq500). Deconvolution was performed on the ICMandTE dataset. Differential expression, with embryo sex and diameter as cofactors and gene set enrichment analysis (GO BP, KEGG, REACTOME databases) were performed using a false discovery rate <0.05 cutoff. Only a few genes were altered (ICM: n=18; TE: n=6) but several gene sets were perturbed (ICM: n=62; TE: n=50) by maternal parity. In YM, only pathways related to transcription, RNA processing and vesicle transport functions were enriched in the ICM whereas only pathways related to RNA localization were enriched in TE. In YN, while only gene sets related to ribosomes and extracellular matrix were enriched in the ICM, functions related to energy and lipid metabolism, lipid transport and interleukin-1 signaling were enriched in the TE. In conclusion, several genes and pathways are affected in embryos collected from nulliparous mares, with different effects on TE and ICM. Embryo development is altered in nulliparous mares, which could partially explain the term phenotype. Whether differences in gene expression result/induce poor embryo-maternal communication remains to be determined.

## 1. Introduction

In mammalian species, including the horse, it is now well established that the periconceptional and gestational maternal environment affect intra and extra-uterine growth and offspring long-term health [1, 2]. These observations fall within the context of the Developmental Origins of Health and Diseases (DOHaD).

In horses, maternal parity defined as the number of gestations that produced a viable fetus (live or stillborn foal), is one of the main factors affecting the foal intra-uterine development. Indeed, foals born to primiparous mares (mares that have not foaled before) are lighter and smaller at birth and remain smaller until 18 months and lighter until 4 month of age compared to controls born to multiparous dams [3–13]. Their insulin sensitivity is higher than that of foals born to multiparous mares, and these data suggest that the normal decrease in insulin sensitivity observed in relation with foal age is delayed [13]. Similarly, testicular maturation is also delayed in foals born to primiparous mares [13]. These alterations in morphology and physiology of foals born to primiparous dams seem to be related to poorer performances in show jumping or on the racecourse than those of subsequent foals born to the same mare [14, 15].

For a long time, these differences in mares’ first born foals have been attributed to the need for a first gestation priming for the uterus to be able to reach its optimal size and vascularisation and fully support feto-placental developmental needs [16]. Indeed, primiparous mares produced lighter and less voluminous placentas than multiparous ones [8,12,13,16]. In horses, placentation is diffuse and the epitheliochorial placenta is in contact with the entire surface of the uterus [17, 18]. Most feto-maternal exchanges occur through branched vascular structures that form interdigitations with the mare endometrium, called microcotyledons, that maximize nutrient exchanges by increasing feto-maternal contact surface [17–19]. Reduced placental volume and weight are associated with reduced foal development in first born foals and suggest that primiparity could be a form of intra-uterine growth restriction in horses.

The placenta derives from the equine embryo trophoblast. Its later efficacy is conditioned by proper implantation and development. Implantation takes place around 35-38 days post ovulation [20]. Prior to that, the equine embryo develops free in the uterus and depends on direct support of uterine secretions for its development. Impaired pre-implantation development in nulliparous mares could play a role in the reduced size of both term placenta and newborn foal. The few existing studies that consider maternal parity on fertility effects are controversial. While some found that parity did not affect fertility [21–27], others reported that mares that have never foaled have reduced embryo and fetal mortality compared to mares that previously foaled [27–33]. Confounding effects of maternal parity and age is probably the source of those discrepancies. Indeed, in a recent epidemiological study considering the effect of parity only in mares older than 10 years, there is a cumulative negative effect of nulliparity and aging on the rates of pregnancy at 14 days post-ovulation (ED and PCP, personal communication). Maternal age have been shown to affect oocyte and embryo developmental capacities (for review [34]) as well as gene expression in Day 8 embryos [35]. At the opposite, only one study considered the effect of maternal parity on preimplantation embryo and showed alterations of the expression of genes related to embryo development and exchanges with the environment were observed [36]. This study, however, only considered mares older than 10 years, in which uterine degenerative changes had probably occur. As maternal age affects embryo gene expression, it is important to consider maternal parity in young mares. At this time, there is no study considering the effect of parity on gene expression of embryos in young mares.

The aim of this study was to determine the effect of maternal nulliparity in young mares on embryo gene expression at the blastocyst stage. Young (5-6 years old) nulliparous and multiparous mares were inseminated with semen of the same stallion. Day-8 blastocysts were collected, measured and bisected to separate the pure trophoblast (TE_part) from the inner cell mass enriched hemi-embryo (ICMandTE). Gene expression was analyzed by RNA-seq in each compartment.

## 2. Materials and methods

### 2.1. Ethics

The experiment was performed at the experimental farm of IFCE (research agreement C1903602 valid until March 22, 2023). The protocol was approved by the local animal care and use committee (“Comité des Utilisateurs de la Station Expérimentale de Chamberet”) and by the regional ethical committee (“Comité Régional d’Ethique pour l’Expérimentation Animale du Limousin”, approved under N° C2EA - 33 in the National Registry of French Ethical Committees for animal experimentation) under protocol number APAFIS#14963-2018050316037888 v2. All experiments were performed in accordance with the European Union Directive 2010/63EU. The authors complied with the ARRIVE guidelines.

### 2.2. Embryo collection

Twenty-one non-nursing mares (mostly French Anglo-Arabian with some Selle Français) aged from 5 to 6 years old were included in this study. Mares were allocated to one of 2 groups according to their parity: nulliparous (YN, n = 10) and multiparous mares (YM, n = 11). Multiparous mares were defined as dams that had already foaled at least once while nulliparous mares were defined as mares that had never foaled before the experiment. During the experimental protocol, mares were managed in one herd in natural pastures 24h/day with free access to water with no nutritional supplementation but for salt blocks. The experiments took place from April 1^st^ to May 3^rd^, 2019. All mares remained healthy during this period. During the experimentation, mare’s withers’ height and weight were measured. Characteristics of all mares and mares that produced an embryo are detailed in Table 1.

**Table 1:** Mares’ characteristics at embryo collection time.

| Characteristics | Nulliparous (YN) |  | Multiparous (YM) |  |
| --- | --- | --- | --- | --- |
|  | Total<br>(n = 10) | With embryo<br>(n = 6) | Total<br>(n = 11) | With embryo<br>(n = 5) |
| Breed | AA n = 7; SF n = 3 | AA n = 5; SF n = 1 | AA n = 9; SF n = 1;<br>SB n=1 | AA |
| Age (*in years*) | 6.00 ± 0.00 | 6.00 ± 0.00 | 6.00 ± 0.00 | 6.00 ± 0.00 |
| Parity (*number of foalings*) | 0 ± 0.00 | 0 ± 0.00 | 1.00 ± 0.00 | 1.00 ± 0.00 |
| Weight (*in kg*) | 535.66 ± 30.72 | 544.55 ± 30.50 | 536.62 ± 44.12 | 524.02 ± 61.39 |
| BCS (*scale 1-5*) | 2.35 ± 0.32 | 2.21 ± 0.25 | 2.16 ± 0.28 | 2.3 ± 0.33 |
| Withers' height (*in cm*) | 159.70 ± 3.33 | 161.67 ± 2.50 | 157.95 ± 3.74 | 157.40 ± 5.41 |

AA: Anglo Arab or Anglo-Arabian type; SF: Selle Français section A or B; SB: Saddlebred. Age, parity,
weight, and height are presented as mean ± SD

Mares were monitored as previously described [35]. Briefly, the mares’ estrous period was monitored routinely by ultrasound with a 5MHz trans-rectal transducer. During estrus, ovulation was induced with a single injection of human chorionic gonadotropin (i.v.; 750 - 1500IU; Chorulon® 5000; MSD Santé animale, France) as soon as one ovarian follicle >35mm in diameter was observed, together with marked uterine edema. Ovulation usually takes place within 48h, with > 80% occurring 25 to 48h after injection [37, 38]. At the same time, mares were inseminated once with fresh or fresh overnight cooled semen containing at least 1 billion motile spermatozoa from a single fertile stallion. Ovulation was confirmed within the next 48 hours by ultrasonography.

Embryos were collected by non-surgical uterine lavage using prewarmed (37°C) lactated Ringer’s solution (B.Braun, France) and EZ-Way Filter (IMV Technologies, France) 10 days after insemination, i.e., approximately 8 days post ovulation. Just after embryo collection, mares were treated with luprotiol an analogue of prostaglandin F2α (i.m; 7.5 mg; Prosolvin, Virbac, France).

The aim of the embryo collection was to obtain 5 embryos/group with each embryo coming from a different mare. Therefore, some mares that failed to produce an embryo at their first attempt were bred again for a second attempt.

### 2.3. Embryo bisection and RNA extraction

Using a binocular magnifying glass, collected embryos were immediately photographed with a size standard to subsequently determine embryo diameter using ImageJ® software (version 1.52a; National Institutes of Health, Bethesda, MD, USA). Embryos were then washed 4 times in commercially available Embryo holding medium (IMV Technologies, France) at 34°C and bisected with a microscalpel under binocular magnifying glass to obtain a trophoblast (TE_part) and an inner cell mass enriched (ICMandTE) hemi-embryo. At this stage, the TE_part is composed of trophectoderm and endoderm whereas the ICM is composed of epiblast layered on the internal side by endoderm cells [39, 40]. Immediately after bisection, RNA extraction of each hemi-embryo was started in extraction buffer (PicoPure RNA isolation kit, Applied Biosystems, France) for 30 min at 42°C prior to storage at −80°C. RNA was extracted later from each hemi-embryo using PicoPure RNA isolation kit (PicoPure RNA isolation kit, Applied Biosystems, France), which included a DNAse treatment, following the manufacturer’s instructions. RNA quality and quantity were assessed with the 2100 Bioanalyzer system using RNA 6000 Pico kit (Agilent Technologies, France) according to the manufacturer’s instructions.

### 2.4. RNA sequencing

Five nanograms of total RNA were mixed with ERCC spike-in mix (Thermofisher Scientific, France) according to manufacturer’s recommendations. Messenger RNAs were reverse transcribed and amplified using the SMART-Seq V4 ultra low input RNA kit (Clontech, France) according to the manufacturer recommendations. Nine PCR cycles were performed for each hemi-embryo. cDNA quality was assessed on an Agilent Bioanalyzer 2100, using an Agilent High Sensitivity DNA Kit (Agilent Technologies, France). Libraries were prepared from 0.15 ng cDNA using the Nextera XT Illumina library preparation kit (Illumina, France). They were pooled in equimolar proportions and sequenced (Paired end 50-34 pb) on NextSeq500 instrument, using a NextSeq 500 High Output 75 cycles kit (Illumina, France). Demultiplexing was performed with bcl2fastq2 version 2.2.18.12 (Illumina, France) and adapters were trimmed with Cutadapt version 1.15 [41]. Only reads longer than 10pb were kept.

### 2.5. RNA mapping and counting

As previously described [35], alignment was performed using STAR version 2.6 [42] on previously modified Ensembl 99 EquCab3.0 assembly and annotation. Genes were then counted with FeatureCounts [43] from Subreads package version 1.6.1.

### 2.6. Data analysis

All statistical analyses were performed by comparing YN to YM (YM set as reference group) using R version 4.0.2 [44] on Rstudio software version 1.3.1056 [45].

Embryo were sexed using *X Inactive Specific Transcript (XIST)* expression as previously described [35]. Six embryos were determined as female (2 in the YN group and 4 in the YM group) while 5 were considered as male (4 in the YN group, and 1 in the YM group).

#### 2.6.1. Embryo recovery and fertility rate, embryo diameter and total RNA content analysis

Embryo recovery rates (ERR) per mare and per ovulation were calculated as the number of attempts with at least one embryo collected/total number of attempts. Both were analyzed using the Exact Fisher test to determine if maternal parity influenced embryo recovery.

For total RNA content analyses, as embryos were bisected without strict equality for each hemi-embryo, a separate analysis of ICMandTE and TE_part RNA quantities would not have been meaningful. Thus, ICMandTE and TE_part RNA quantities were summed up. RNA quantity and embryo diameter were analyzed using a linear model of nlme package version 3.1-148 [46] including maternal parity and embryo sex, followed by 1000 permutations using PermTest function from pgirmess package version 1.6.9 [47]. Variables were kept in the subsequent models when statistically significant differences were observed. Differences were considered as significant for p < 0.05.

#### 2.6.2. Deconvolution of gene expression in ICMandTE using DeMixT

The deconvolution method has already been described in equine embryos [35]. Briefly, this method enables the estimation of the relative gene expression of TE and ICM cell types within the hemi-embryo ICMandTE which is composed of both trophoblast and inner cell mass in unknown relative proportions. After filtering all genes with 3 non-null count values in at least one group (YN or YM) per hemi-embryo (ICMandTE or TE_part), removing genes with a null variance in TE_part and adding the value “1” to all count values in ICMandTE and TE_part datasets, deconvolution was performed using the DeMixT R package version 1.4.0 [48, 49]. Output datasets were DeMixT_ICM_cells and DeMixT_TE_cells, corresponding to the deconvoluted gene expression in ICM cells and TE cells of ICMandTE, respectively.

At the end of deconvolution, a quality check was automatically performed by the DeMixT R package with the TE_part used as reference for DeMixT_TE_cells. Genes were automatically filtered out if the difference between average deconvoluted expression of reference cells in mixed samples and average expression of reference cells > 4.

Outputs of DeMixT_ICM_cells *vs* DeMixT_TE_cells, DeMixT_ICM_cells *vs* TE_part and ICMandTE *vs* TE_part were compared with Deseq2 version 1.28.1 [50] to confirm that the deconvolution was effective at separating gene expression. To check if deconvolution was efficient, as previously described [35], the expression of several genes proper to ICM and TE cells in equine embryos identified using literature search [51] was compared before and after deconvolution. Results of these analyses were represented through manually drawn Venn diagrams as well as principal component analysis graphics of individuals, using ggplot2 version 3.3.3 [52] and factoextra version 1.0.7 [53].

#### 2.6.3. Maternal parity comparison for gene expression

All genes with an average expression <10 counts in both YN and YM per hemi-embryo (ICM or TE) were filtered out on the DeMixT_ICM_cells and TE_part datasets. Differential analyses were performed with Deseq2 version 1.28.1 [50] with the YM group as reference, without independent filtering. Genes were considered differentially expressed (DEG) for FDR <0.05 after Benjamini-Hochberg correction (also known as false discovery rate, FDR). As ovulation was checked only every 48h and because embryos growth is exponential in the uterus, embryo diameter was considered as a cofactor in the model as well as embryo sex.

Equine Ensembl IDs were converted into Human Ensembl IDs and Entrez Gene names using gorth function in gprofiler2 package version 0.1.9 [54]. Genes without Entrez Gene names using gprofiler2 were manually converted when Entrez Gene names were available, using Ensembl web search function [55]. GO molecular function and GO Biological process annotations of genes were obtained from Uniprot website.

#### 2.6.4. Gene set enrichment analyses (GSEA)

After log transformation using RLOG function of DESeq2 version 1.28.1, gene set enrichment analyses (GSEA) were performed on expressed genes using GSEA software version 4.0.3 (Broad Institute, Inc., Massachusetts Institute of Technology, and Regents of the University of California) [56, 57] to identify biological gene sets disturbed by maternal parity. Molecular Signatures Databases [58] version 7.1 (C2: KEGG: Kyoto Encyclopedia of Genes and Genomes; REACTOME, C5: BP: GO biological process) were used to identify most perturbed pathways. Pathways were considered significantly enriched for FDR< 0.05. When the normalized enrichment score (NES) was positive, the gene set was enriched in the YN group while when NES was negative, the gene set was enriched in the YM group.

If applicable, as previously described in equine embryos [35], enriched terms from GO BP, KEGG and REACTOME databases were represented using SUMER analysis from SUMER R package version 1.1.5 and using FDR q-values [59]. Results were represented with graphs modified using Cytoscape version 3.8.2 [60]. In these graphs, gene sets are represented by nodes and the gene set size is represented by the size of the node. Node shape represents the gene set database (GO BP, KEGG or REACTOME). Blue nodes represent gene sets enriched in YN (NES > 0) while green nodes represent gene sets enriched in YM (NES < 0). Edge width represents the level of connection between representative gene sets (thinner edges represent the first clustering while thicker edges represent the second clustering of the affinity propagation algorithm).

As SUMER is not able to consider only genes that participate to enrichment in GSEA, pathways with genes in common are grouped together, although genes in common are not the ones that participate to the enrichment. To better understand groups, therefore, authors, first recovered genes that were enriched in each pathway of a common group of gene sets according to SUMER analysis. Then, they only considered genes in common between pathways in one group to better qualify the function that was altered by maternal parity.

## 3. Results

### 3.1. Embryo recovery rates, diameter, total RNA content and quality and progesterone concentrations

Altogether, 25 embryo collections were performed (13 in YN and 12 in YM, 4 mares being flushed twice) and 12 embryos were obtained (7 from 6 YN mares and 5 from 5 YM mares). One young nulliparous mare produced twin embryos.

Embryo recovery rate per mare was 46% and 42% in YN and YM, respectively and did not differ between groups (p = 1).

Altogether, 1 and 2 double ovulations were observed, respectively, in YN and YM. The embryo recovery rate per ovulation at the time of embryo collection was not different according to group (50% in YN and 36% in YM, p = 0.70).

All embryos were expanded blastocysts grade I or II according to the embryo classification of McKinnon and Squires [61]. For the twin collection, embryos diameters were 580µm and 591µm. As only one embryo per mare was required, the 580µm diameter was randomly chosen for further analysis. Altogether, only 6 YN and all 5 YM embryos collected were RNA sequenced. Embryo diameter ranged from 457µm to 2643µm, with no effect of group on embryo diameter (p = 0.18). In embryos selected for RNA sequencing, there was no effect of embryo sex on its size (p = 0.63). RNA yield per embryo ranged from 12.0 ng to 2915.5 ng and was not related to parity (p = 0.07) nor embryo sex (p = 0.77).

The median RNA Integrity Number (RIN) was 9.6 (8.9 - 10 range). Between 39.7 and 69.5 million reads per sample were obtained after trimming. On average, 70.94% of the reads were mapped on the modified EquCab 3.0 using STAR and 66.45% were assigned to genes by featureCounts.

### 3.2. Deconvolution of gene expression to discriminate ICM and TE gene expression in ICMandTE hemi-embryos

After selecting genes with more than 3 non null count values in at least one group (YN or YM) per hemi-embryo (ICMandTE or TE_part), 16,901 genes were conserved for deconvolution. In addition, 67 genes were removed because their variance was null in the TE_part. For these genes, the mean count in ICMandTE samples was above 110 counts. The deconvolution quality of all gene was sufficient. Therefore, at the end of the deconvolution algorithm, 16,834 genes were available for differential analysis.

Before deconvolution, 681 genes were differentially expressed (FDR < 0.05) between the ICMandTE and the TE_part (Fig. 1a). After deconvolution, the comparison between DeMixT_ICM_cells and DeMixT_TE_cells yielded 6,171 differentially expressed genes while the comparison DeMixT_ICM_cells *vs* TE_part yielded 5,262 differentially expressed genes, with 4713 genes in common with the previous comparison (70%). Moreover, 677 of the initially 681 differentially expressed genes before deconvolution were also identified as differentially expressed in both post-deconvolution analyses. Only in the comparison DeMixT_ICM_cells *vs* TE_part, 3 among the 4 remaining genes were identified. On the PCA graph of individuals, ICMandTE and TE_part were partly overlapping (Fig 1b). DeMixT_TE_cells and TE_part superposed well, suggesting that datasets before and after deconvolution have a similar global gene expression; whereas the DeMixT_ICM_cells group is clearly separated from others on Axis 1 (22.3% of variance), indicating that the deconvolution effectively enabled the separation of gene expression of the two cell types in the mixed part (ICMandTE).

**Fig. 1:**
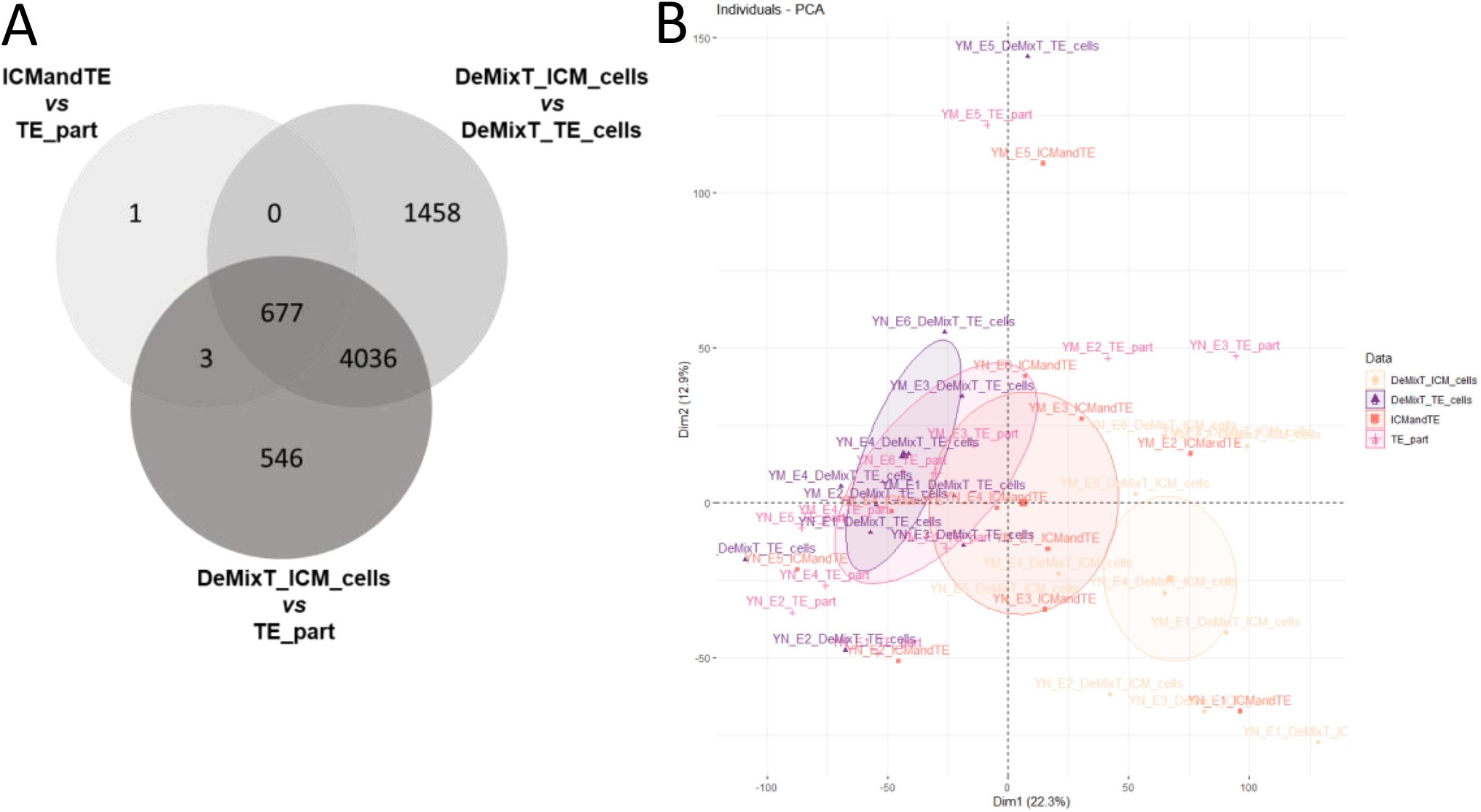
Gene expression in ICM and TE before and after deconvolution using DeMixT. a) Venn diagram of genes differentially expressed in the different analyses: ICMandTE *vs* TE_part (before deconvolution), DeMixT_ICM_cells *vs* DeMixT_TE_cells (after deconvolution) and DeMixT_ICM_cells *vs* TE_part (gene expression of ICM after deconvolution *vs* gene expression in TE_part without deconvolution); b) Principal Component Analysis of gene expression from DeMixT_ICM_cells, DeMixT_TE_cells, ICMandTE and TE part datasets. Deconvolution was used to isolate gene expression of ICM and TE cells in ICMandTE hemi-embryos. ICMandTE: inner cell mass + trophoblast; TE_part: pure trophoblast. Here trophoblast represents trophectoderm + endoderm.

On the 12 genes previously identified by Iqbal et al. as more expressed in the ICM [51], one had to be removed before deconvolution because its variance in the TE was zero (*ENSECAG00000010653*, annotated as SRY-Box Transcription Factor 2, *SOX2*). On the 11 remaining genes, 4 were also more expressed in the ICMandTE *vs* TE_part comparison (Table 2). After deconvolution (comparison DeMixT_ICM_cells *vs* TE_part), 10 out of 11 of these genes were effectively more expressed in the ICM. Iqbal et al. identified 7 genes that were more expressed in the TE. One of those genes was differentially expressed in the comparison ICMandTE *vs* TE_part, *i.e.*, before deconvolution. After deconvolution, the expression of 3 of the 7 reported genes, different from the only gene identified before deconvolution, were increased in the TE_part compared to the DeMixT_ICM_cells.

**Table 2:**
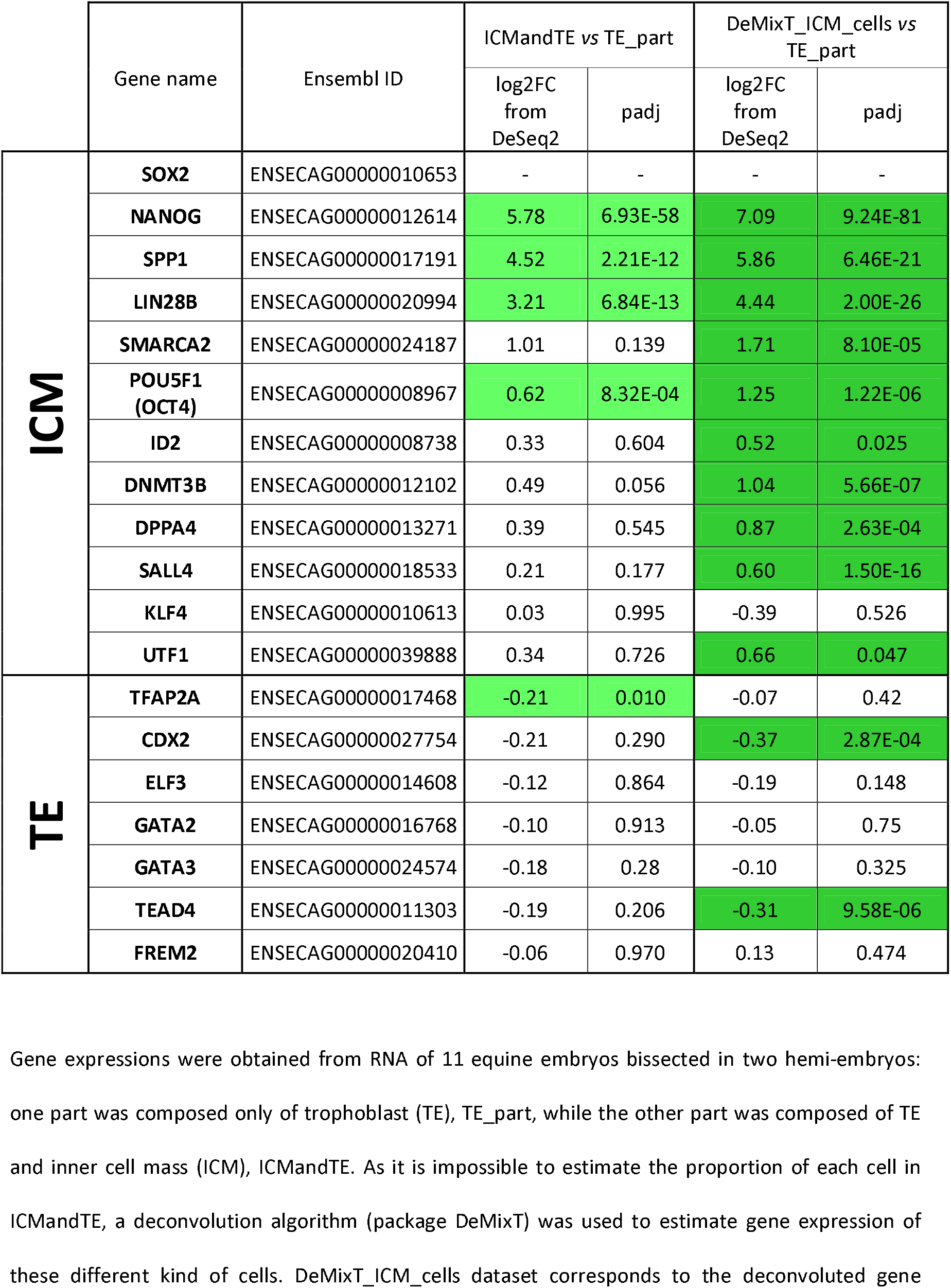
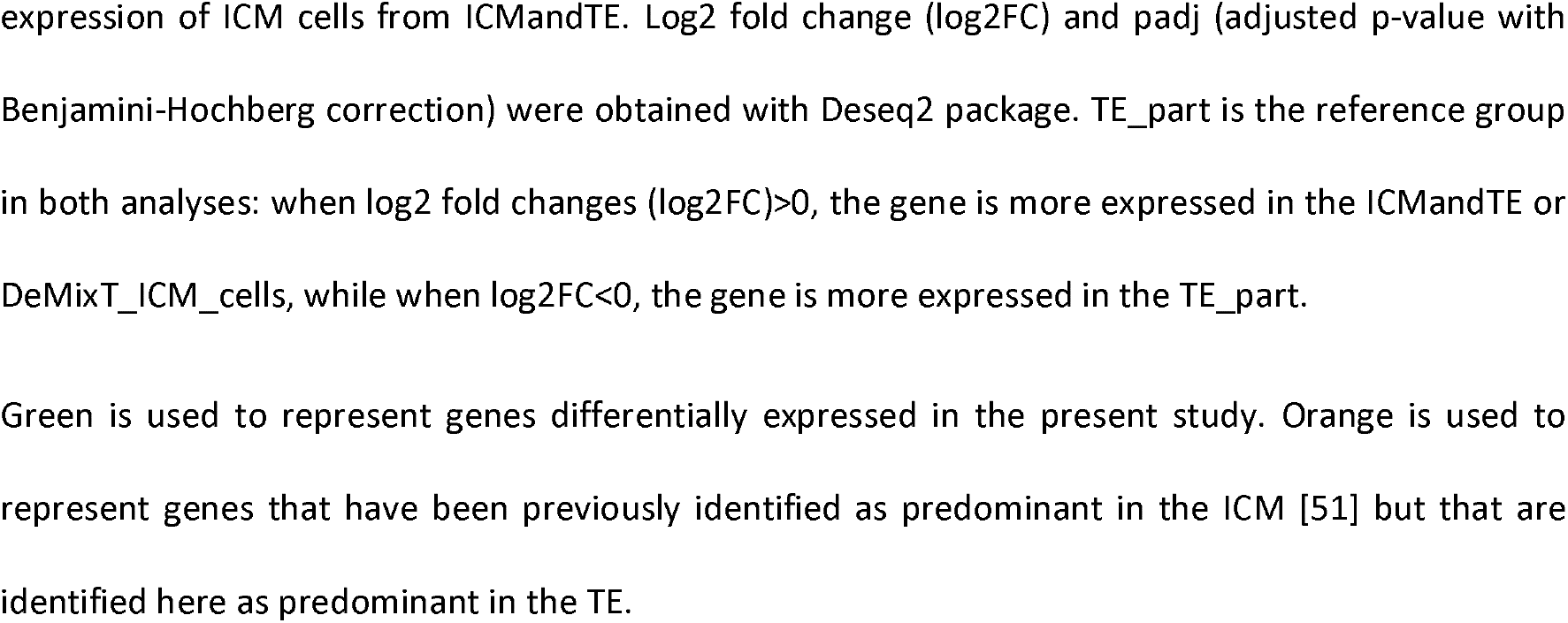
Comparison of the expression of selected genes previously identified as specific to TE or ICM in equine embryos [51], before and after deconvolution

|  | Gene name | Ensembl ID | ICMandTE vs TE_part |  | DeMixT_ICM_cells vs TE_part |  |
| --- | --- | --- | --- | --- | --- | --- |
|  |  |  | log2FC from DeSeq2 | padj | log2FC from DeSeq2 | padj |
| **ICM** | **SOX2** | ENSECAG00000010653 | - | - | - | - |
|  | **NANOG** | ENSECAG00000012614 | 5.78 | 6.93E-58 | 7.09 | 9.24E-81 |
|  | **SPP1** | ENSECAG00000017191 | 4.52 | 2.21E-12 | 5.86 | 6.46E-21 |
|  | **LIN28B** | ENSECAG00000020994 | 3.21 | 6.84E-13 | 4.44 | 2.00E-26 |
|  | **SMARCA2** | ENSECAG00000024187 | 1.01 | 0.139 | 1.71 | 8.10E-05 |
|  | **POU5F1 (OCT4)** | ENSECAG00000008967 | 0.62 | 8.32E-04 | 1.25 | 1.22E-06 |
|  | **ID2** | ENSECAG00000008738 | 0.33 | 0.604 | 0.52 | 0.025 |
|  | **DNMT3B** | ENSECAG00000012102 | 0.49 | 0.056 | 1.04 | 5.66E-07 |
|  | **DPPA4** | ENSECAG00000013271 | 0.39 | 0.545 | 0.87 | 2.63E-04 |
|  | **SALL4** | ENSECAG00000018533 | 0.21 | 0.177 | 0.60 | 1.50E-16 |
|  | **KLF4** | ENSECAG00000010613 | 0.03 | 0.995 | -0.39 | 0.526 |
|  | **UTF1** | ENSECAG00000039888 | 0.34 | 0.726 | 0.66 | 0.047 |
| **TE** | **TFAP2A** | ENSECAG00000017468 | -0.21 | 0.010 | -0.07 | 0.42 |
|  | **CDX2** | ENSECAG00000027754 | -0.21 | 0.290 | -0.37 | 2.87E-04 |
|  | **ELF3** | ENSECAG00000014608 | -0.12 | 0.864 | -0.19 | 0.148 |
|  | **GATA2** | ENSECAG00000016768 | -0.10 | 0.913 | -0.05 | 0.75 |
|  | **GATA3** | ENSECAG00000024574 | -0.18 | 0.28 | -0.10 | 0.325 |
|  | **TEAD4** | ENSECAG00000011303 | -0.19 | 0.206 | -0.31 | 9.58E-06 |
|  | **FREM2** | ENSECAG00000020410 | -0.06 | 0.970 | 0.13 | 0.474 |

Gene expressions were obtained from RNA of 11 equine embryos bisected in two hemi-embryos: one part was composed only of trophoblast (TE), TE\_part, while the other part was composed of TE and inner cell mass (ICM), ICMandTE. As it is impossible to estimate the proportion of each cell in ICMandTE, a deconvolution algorithm (package DeMixT) was used to estimate gene expression of these different kind of cells. DeMixT\_ICM\_cells dataset corresponds to the deconvoluted gene
expression of ICM cells from ICMandTE. Log2 fold change (log2FC) and padj (adjusted p-value with Benjamini-Hochberg correction) were obtained with Deseq2 package. TE\_part is the reference group in both analyses: when log2 fold changes (log2FC)>0, the gene is more expressed in the ICMandTE or DeMixT\_ICM\_cells, while when log2FC<0, the gene is more expressed in the TE\_part.
Green is used to represent genes differentially expressed in the present study. Orange is used to represent genes that have been previously identified as predominant in the ICM [51] but that are identified here as predominant in the TE.

All of these results validate the deconvolution procedure and justify the use of data from the DeMixT_ICM_cells file. In the following results, the TE_part was used as representative of TE and DeMixT_ICM_cells was used as representative of gene expression in the ICM.

### 3.3. Sample selection

One embryo (YM) was larger than 2000 µm while all other embryos were smaller than 1400µm in diameter (Supplementary Figure 1). Embryo size has been shown to affect equine embryo gene expression [62]. Thus, the analysis was performed both with or without this large embryo to check if results were affected. All but one differential expressed genes identified with the largest embryo were also found differentially expressed without it (Supplementary Figure 2). Nevertheless, to limit size effect, the analyses described below are those without the largest embryo, where only 6 YN and 4 YM embryos were analyzed. The results of the differential analyses that were performed including the 2643µm large YM embryo are shown in Supplementary Tables 1 and 2.

### 3.4. Differential gene expression in deconvoluted ICM cells

After retaining only genes with an average expression ≥ 10 counts in at least one maternal parity group andhemi-embryo, 14,418 genes were considered as expressed in the YN or YM embryos ICM cells. Only 18 genes were differentially expressed (12 downregulated and 6 upregulated in YN) (Fig. 2 and Supplementary table 3). Respectively, 11 and 5 genes out of the down- and upregulated genes were associated to a protein known and described in human. These 16 genes an gene sets determined from Uniprot in wich they are susceptible to play a role are presented in Table 3.

**Fig. 2:**
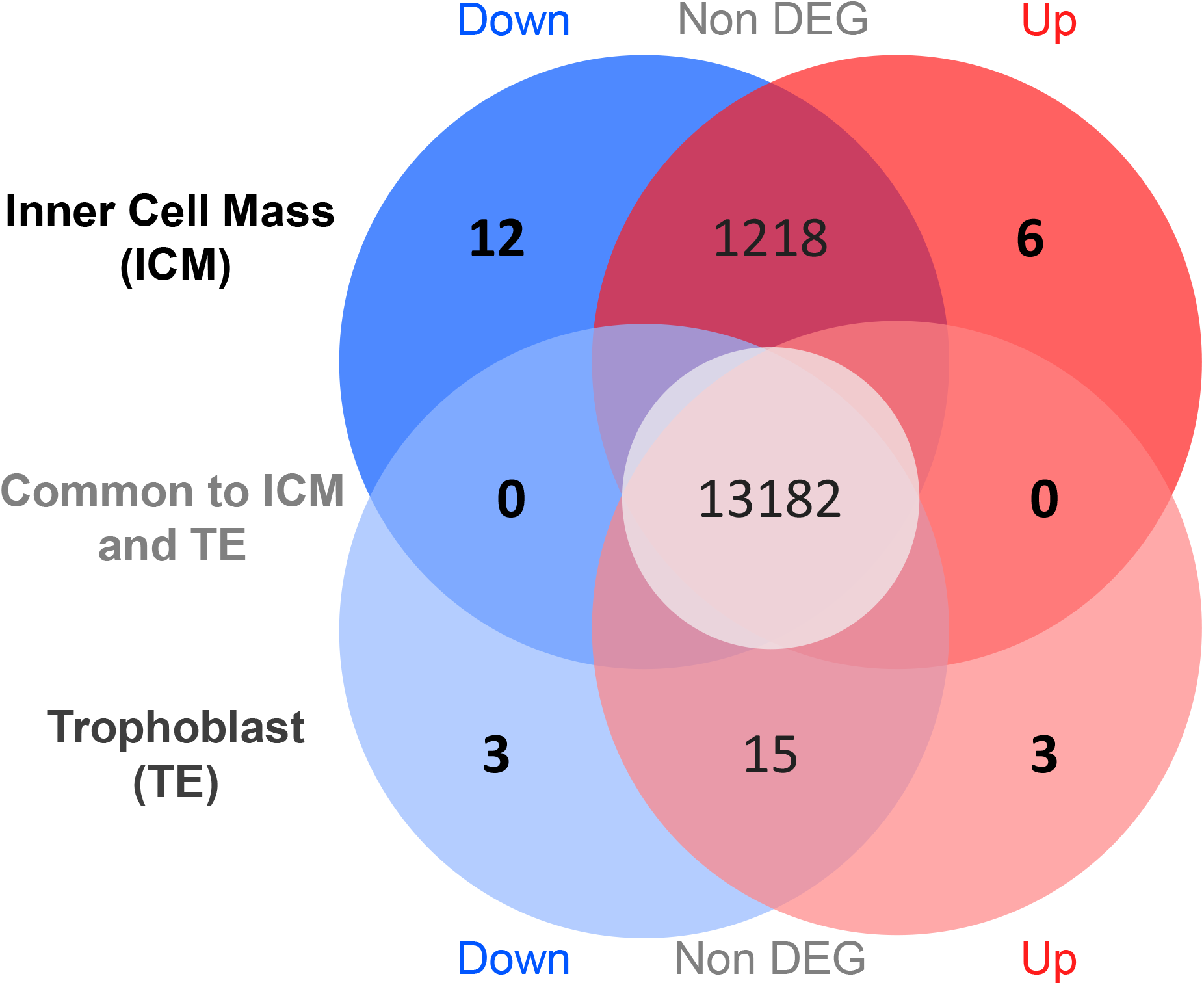
Analysis of differentially expressed genes (DEG) in embryos according to maternal parity. A) representation of down- (blue) and up- (red) regulated DEG in ICM (from DeMixT_ICM_cells data obtained after deconvolution of ICMandTE using DeMixT R package [48, 49]) and TE (from TE_part dataset) of embryos from ON *vs* OM. DEG: Differentially Expressed Genes (FDR < 0.05); TE: Trophoblast; ICM: Inner Cell Mass; ON: Old nulliparous mares; OM: Old multiparous mares

**Table 3:** Up- and down-regulated genes coding for a protein in the inner cell mass of equine embryos according to mare parity

| Ensembl Name | Entrez Gene ID | Description | GO Molecular function | GO Biological Process | log2 Fold Change | padj |
| --- | --- | --- | --- | --- | --- | --- |
| ENSECAG00000029895 | *MAGEB16* | MAGE family member B16 | Tumor antigen |  | -3.24 | 0.013 |
| ENSECAG00000017619 | *NOXO1* | NADPH oxidase organizer 1 | Phospholipid binding<br>Superoxide-generating NADPH oxidase activator activity | Extracellular matrix disassembly<br>Positive regulation of cell killing<br>Regulation of hydrogen peroxide metabolic process<br>Superoxide metabolic process<br>Regulation of respiratory burst | -2.67 | 0.034 |
| ENSECAG00000023392 | *ZBTB8A* | zinc finger and BTB domain containing 8A | Metal ion binding<br>DNA binding | Regulation of transcription by RNA polymerase II<br>Transcription regulation | -2.02 | 5.84E-05 |
| ENSECAG00000019702 | *VPS52* | VPS52 subunit of GARP complex | Syntaxin binding | Ectodermal cell differentiation<br>Embryonic ectodermal digestive tract development<br>Endocytic recycling<br>Lysosomal transport<br>Protein transport | -1.26 | 0.001 |
| ENSECAG00000011960 | *DESI1* | desumoylating isopeptidase 1 | Hydrolase<br>Identical protein binding | Protein desumoylation<br>Protein export from nucleus<br>Protein modification by small protein removal | -1.23 | 0.001 |
| ENSECAG00000020433 | *TENM3* | teneurin transmembrane protein 3 | Cell adhesion molecule binding<br>Protein heterodimerization activity<br>Protein homodimerization activity | Cell adhesion<br>Differentiation<br>Neuron development<br>Signal transduction | -1.00 | 0.015 |
| ENSECAG00000015867 | *PAQR3* | progesterone and<br>adipoQ receptor<br>family member 3 | Signaling receptor activity | Negative regulation of MAP kinase<br>activity<br>Negative regulation of neuron<br>projection development<br>Negative regulation of protein<br>phosphorylation | -0.84 | 0.018 |
| ENSECAG00000004931 | *MARS2* | methionyl-tRNA<br>synthetase 2,<br>mitochondrial | Aminoacyl-tRNA synthetase<br>Ligase<br>ATP binding | Protein biosynthesis<br>Methionyl-tRNA aminoacylation<br>tRNA aminoacylation for protein<br>translation | -0.82 | 0.020 |
| ENSECAG000000034815 | *VHL* | Von Hippel-Lindau<br>tumor suppressor | Enzyme binding<br>Transcription factor binding | Ubl conjugation pathway<br>Cell morphogenesis<br>Negative regulation of apoptotic<br>process<br>Negative regulation of cell population<br>proliferation<br>Negative regulation of gene<br>expression<br>Negative regulation of transcription<br>by RNA polymerase II<br>Positive regulation of cell<br>differentiation<br>Positive regulation of transcription<br>Protein stabilization<br>Proteolysis | -0.81 | 0.018 |
| ENSECAG000000007262 | *EEA1* | early endosome<br>antigen 1 | GTP-dependant protein binding<br>Protein homodimerization activity | Early endosome to late endosome<br>transport<br>Endocytosis<br>Vesicle fusion | -0.79 | 0.015 |
| ENSECAG000000009590 | *GSTCD* | glutathione S-transferase C-terminal domain<br>containing |  |  | -0.72 | 0.037 |
| ENSECAG00000014243 | *GATM* | glycine<br>amidinotransferase | Amidinotransferase activity | Creatine metabolic process<br>Multicellular organism development | 0.73 | 0.018 |
| ENSECAG00000041817 | *RBP1* | retinol binding<br>protein 1 | Retinoid binding | Lipid homeostasis<br>Retinoic acid biosynthetic process | 1.13 | 0.020 |
| ENSECAG00000010447 | *EFEMP1* | EGF containing<br>fibulin extracellular<br>matrix protein 1 | Calcium ion binding<br>Epidermal growth factor receptor<br>binding | Embryonic eye morphogenesis<br>Epidermal growth factor receptor<br>signaling pathway<br>Regulation of transcription, DNA-templated | 1.95 | 0.008 |
| ENSECAG00000010385 | *MET* | MET proto-<br>oncogene,<br>receptor tyrosine<br>kinase | ATP binding<br>Protein phosphatase binding<br>Transmembrane receptor protein<br>tyrosine kinase activity | Branching morphogenesis of an<br>epithelial tube<br>Cell migration<br>Cell surface receptor signaling<br>pathway<br>MAPK cascade<br>Phagocytosis<br>Positive regulation of microtubule<br>polymerization<br>Positive regulation of transcription by<br>RNA polymerase II | 2.83 | 0.013 |
| ENSECAG00000022277 | *MAOA* | Monoamine<br>oxidase A | Monoamine oxidase activity<br>Oxidoreductase | Cellular biogenic amine metabolic<br>process<br>Cytokine-mediated signaling pathway<br>Dopamine catabolic pathway<br>Positive regulation of signal<br>transduction | 4.51 | 0.001 |

8 Log2 Fold-Change<0 indicates down-regulation of the gene in embryos from nulliparous mares, also indicated in green; Log2 Fold-Change>0 indicates up-regulation of the 9 gene in embryos from nulliparous mares, also indicated in blue.

### 3.5. Differential gene expression in the TE part

In the TE, 13,203 genes were considered as expressed in YN or YM. Only 6 were differentially expressed (Supplementary table 4) with half being down and up-regulated in YN (Fig. 2). Except one that was a long noncoding RNA, all other genes were associated to a known protein in human. These genes are presented in Table 4 with the pathways in which they are susceptible to play a role.

**Table 4:**
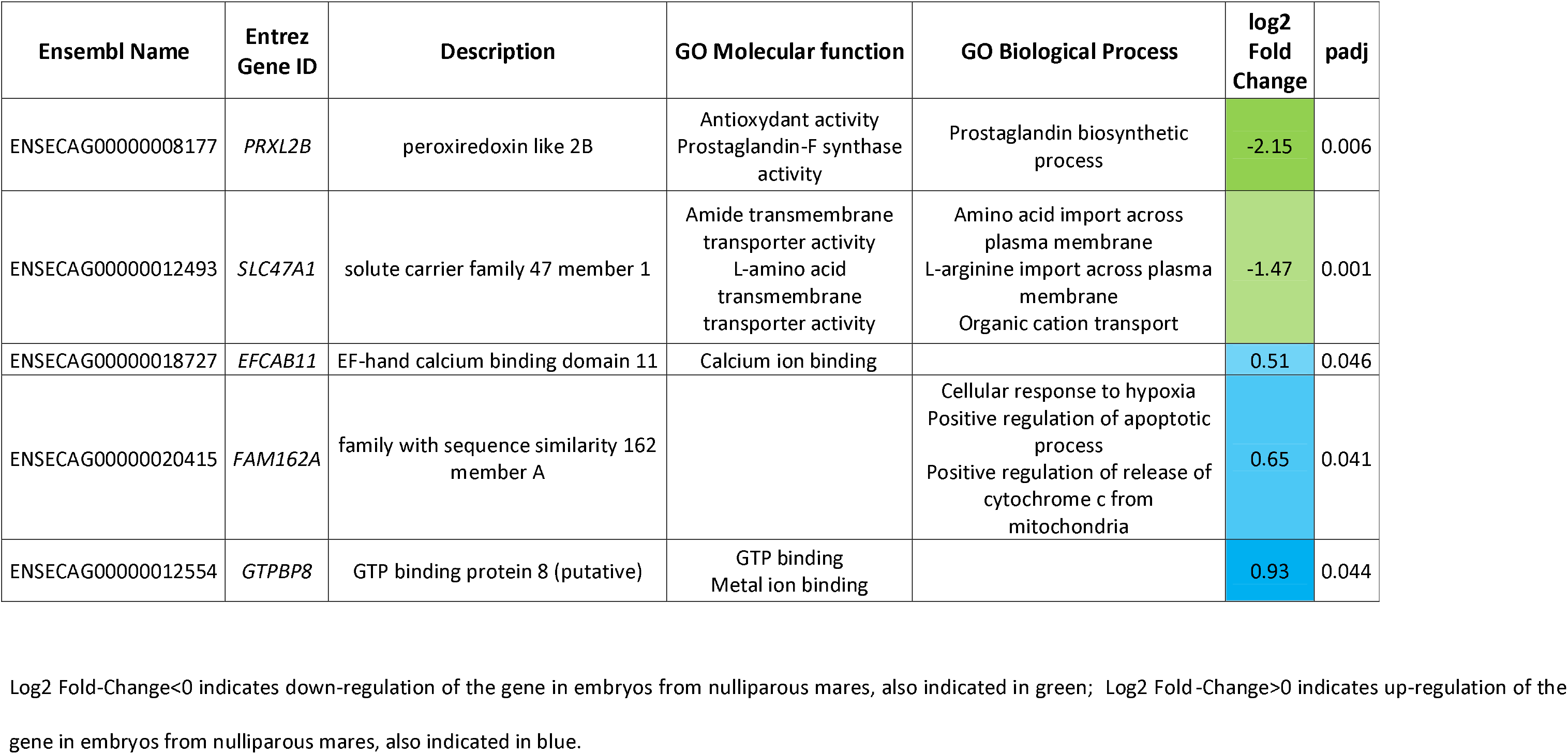
Up- and down-regulated genes coding for a protein in the trophoblast part of equine embryos according to mare parity

| Ensembl Name | Entrez Gene ID | Description | GO Molecular function | GO Biological Process | log2 Fold Change | padj |
| --- | --- | --- | --- | --- | --- | --- |
| ENSECAG00000008177 | *PRXL2B* | peroxiredoxin like 2B | Antioxydant activity<br>Prostaglandin-F synthase activity | Prostaglandin biosynthetic process | -2.15 | 0.006 |
| ENSECAG00000012493 | *SLC47A1* | solute carrier family 47 member 1 | Amide transmembrane transporter activity<br>L-amino acid transmembrane transporter activity | Amino acid import across plasma membrane<br>L-arginine import across plasma membrane<br>Organic cation transport | -1.47 | 0.001 |
| ENSECAG00000018727 | *EFCAB11* | EF-hand calcium binding domain 11 | Calcium ion binding |  | 0.51 | 0.046 |
| ENSECAG00000020415 | *FAM162A* | family with sequence similarity 162 member A |  | Cellular response to hypoxia<br>Positive regulation of apoptotic process<br>Positive regulation of release of cytochrome c from mitochondria | 0.65 | 0.041 |
| ENSECAG00000012554 | *GTPBP8* | GTP binding protein 8 (putative) | GTP binding<br>Metal ion binding |  | 0.93 | 0.044 |

2 Log2 Fold-Change<0 indicates down-regulation of the gene in embryos from nulliparous mares, also indicated in green; Log2 Fold-Change>0 indicates up-regulation of the 3 gene in embryos from nulliparous mares, also indicated in blue.

### 3.6. Gene set enrichment analysis in deconvoluted ICM cells

After Entrez Gene ID conversion, 12,892 genes were considered expressed in ICM cells. Fifty-eight GO Biological Process and 4 KEGG pathways were disturbed by maternal parity in ICM cells (Supplementary table 5). After SUMER analysis, 2 and 27 gene sets, respectively enriched in YN and YM, were represented (Fig. 3). They were clustered in 8 groups. The group enriched in YM and clustered under the term “DNA recombination” was composed of genes related to the maintenance of DNA integrity, chromosome segregation and recombination. Enriched in YM groups “NCRNA metabolic process” and ‘Peptidyl lysine trimethylation” contained both, genes related to methylation and transcription. The only gene set enriched in YN in the group “NCRNA metabolic process” was mainly enriched by genes encoding for a subunit of ribosomes that were common with other gene sets enriched in YM. Genes related to ribosomes were, however, not participating in gene set enrichment of other pathways in this cluster. “multi organism localization”, “RNA splicing” and “RNA localization” clusters were composed of genes involved in RNA maturation and transport. The last group enriched in YM was clustered under the term “vesicle targeting” and was containing genes related to intracellular transport. In this group, the pathway “Golgi vesicle transport” included a DEG that was Vacuolar Protein Sorting-Associated Protein 52 Homolog (*VPS52*), up-regulated in the ICM of embryos from YM mares. The only one group enriched in YN was composed of one pathway named “ECM receptor interaction” in which genes related to extracellular matrix (ECM) were observed.

**Fig. 3:**
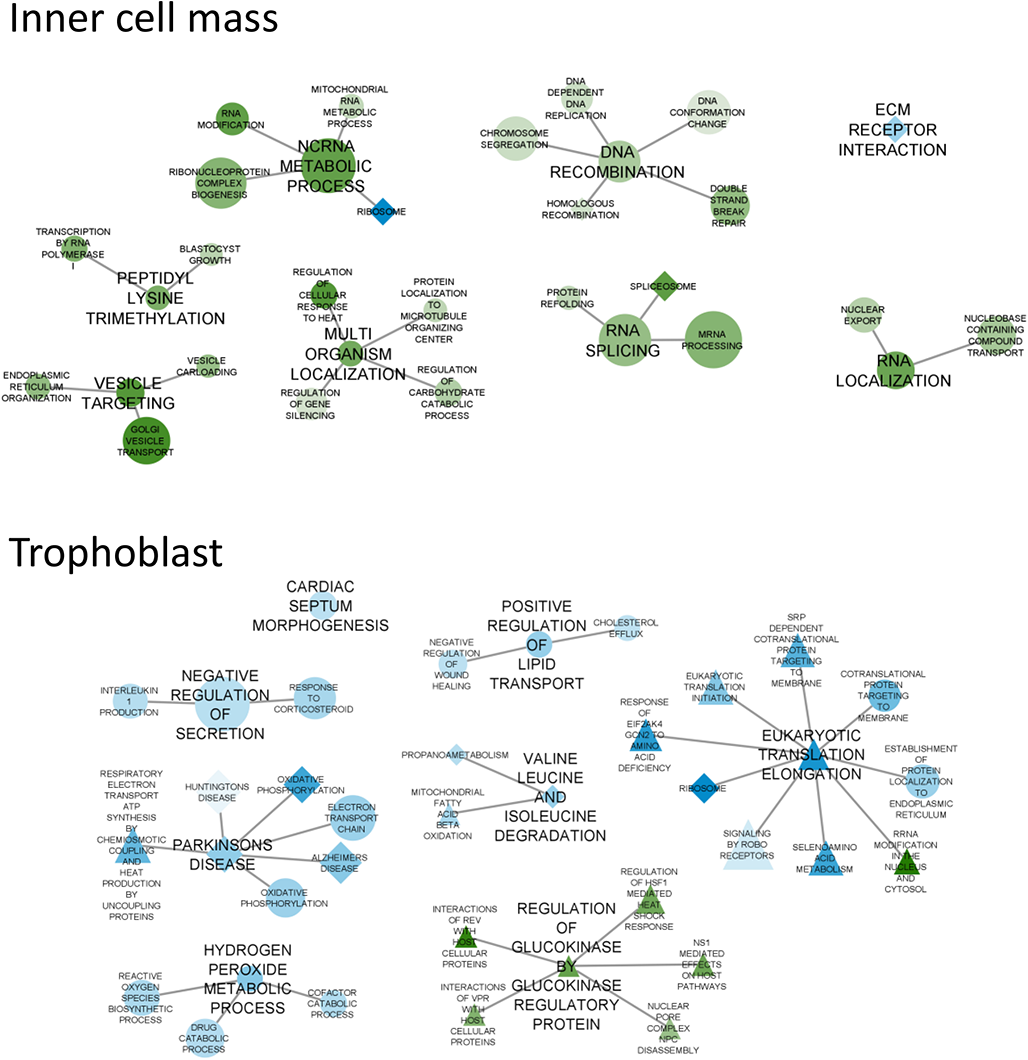
SUMER clustering of GSEA terms clustering of the most perturbed terms in the ICM and TE of embryos according to mares’ parity. Nodes represent altered gene sets in the ICM and TE (FDR <0.05). Node size represents the gene set size. Node shape represents the gene set database: GO BP (circle) or KEGG (diamond) or REACTOME (square). Gene sets are represented in blue if enriched (NES >0) in young nulliparous mares’ embryos and in green if enriched (NES <0) in young multiparous mares’ embryos. The lighter the color, the more the NES is close to 0. Edges represent the level of connection between representative gene sets. This graph was performed using SUMER R package [59] and modified using cytoscape 3.8.2 [60] GSEA: Gene set enrichment analysis; ICM: Inner cell mass; TE: trophoblast; FDR: False Discovery Rate; GO BP: Gene Ontology Biological Process; Kyoto Encyclopedia of Genes and Genomes; NES: Normalized Enrichment Score

### 3.7. Gene set enrichment analysis in TE

After Entrez Gene ID conversion, 11,889 genes were considered expressed in TE from YN or YM embryos. Altogether, 50 gene sets from GO BP, KEGG and REACTOME were perturbed (23 GO BP, 7 KEGG and 20 REACTOME) by maternal parity in young mares (Supplementary table 6). After SUMER analysis, 36 gene sets were represented (Fig. 3) and were clustered in 8 groups. Among them, 7 were enriched in YN. The first group was composed of one gene set named “cardiac septum morphogenesis”. Most genes that participated to the enrichment of this pathway were related to transcriptional factors. The second group was composed of 3 pathways and clustered under the term “negative regulation of secretion”. Genes that participated most to the enrichment of these pathways were related to the innate immunity, more particularly to the production and transport of the interleukin 1 beta (IL1B). Altogether, 3 groups were related to the production of energy inside the cell. Indeed, the cluster under the term “Parkinsons disease” was actually composed of genes with an enriched expression that were related to oxidative phosphorylation. One group was “Hydrogen peroxide metabolic process” where genes that participated the most to the enrichment were involved in the degradation of hydrogen peroxide. The last group involved in energy production was clustered under the term “valine, leucine and isoleucine degradation”. Genes that participated the most to the enrichment of these pathways were directly involved in the beta oxidation of fatty acid. The group under the term “positive regulation of lipid transport” was composed of genes related to the regulation of the transport of lipids and cholesterol. The last group enriched in YN was clustered under the term “eukaryotic translation elongation” and was mostly enriched due to components of the ribosomes that mostly participate in these enrichments. Moreover, the REACTOME pathway “RRNA modification in the nucleus and cytosol” was clustered with this group because of genes that encoded for ribosome components. These genes were, however, not enriched in this particular pathway. Genes that participate to its enrichment, nevertheless, were related to ribosome biogenesis. The only one group enriched in YM was represented by the term “Regulation of glucokinase by glucokinase regulatory protein”. These pathways were mostly enriched in YM because of genes that encode for nucleoporin subunits.

## 4. Discussion

Maternal parity in young mares slightly affected both ICM and TE gene expression without affecting embryo recovery rates nor growth. Although only a few genes were affected by maternal parity, up regulated genes in the ICM of embryos from young nulliparous were involved in lipid, amine and creatinine metabolism, positive regulation of transcription, growth factor signaling and morphogenesis while down regulated were related to extracellular matrix (ECM) disassembly, reactive oxygen species (ROS) metabolism, transcription regulation, endocytosis, protein transport, protein metabolism, MAP kinase signalization and cell differentiation. In the TE, only five known genes were observed differentially expressed. One of the 3 up regulated genes in the TE of embryos from YN was involved in the cell response to hypoxia while the 2 others encode for ion binding proteins. Down regulated genes in the TE of embryos from YN mares were related to prostaglandin metabolism and amino acid/creatinine exchange. Interestingly, while gene set enrichment analysis showed almost only pathway enrichment in YM in the ICM, in the TE, almost all pathways were enriched in YN embryos. In the ICM, after SUMER analysis, the pathways enriched in YM were related to DNA modification, RNA production and maturation and cell transport while gene sets related to the extracellular matrix function and ribosome were enriched in ICM of YN embryos. In the TE, gene sets enriched in embryos from YN were related to immunity, growth factor signaling, phosphorylation oxidative, metabolism of reactive oxygen species, beta oxidation and transport of lipids and ribosome while gene sets enriched in embryos from YM were related to nucleoporins.

In the ICM of embryos from YM mares, enriched gene sets were mostly related to DNA conformation and methylation changes as well as RNA formation, transport and maturation. These results suggest that transcription and regulating pathways are less active in embryos from nulliparous mares. These results are comforted by the fact that teneurin transmembrane protein 3 (*TENM3*) is downregulated in embryos from nulliparous mares. This gene is part of the teneurin family, which encodes for transmembrane proteins that are essential for embryo morphogenesis and nervous system development. The knockdown of these genes in mice and drosophila leads to embryo lethality (for review [63]). Altogether, these results suggested that ICM growth and development would be poorer in embryo from nulliparous compared to multiparous mares but, here, no difference in embryo size had been observed. One hypothesis could be because only ICM seemed affected or because, embryo size at a same age is highly variable as shown in several studies [64–68], although ovulation check was performed twice daily [69]. This huge variation could hide size differences in studies.

In the TE, the gene named “family with sequence similarity 162 member A” (*FAM162A*), also known as E2-Induced Gene 5 Protein (*E2IG5*) or growth and transformation-dependent protein (*HGTD-P*) was up-regulated in embryos from nulliparous mares. This gene is one of the hypoxia inducible factors (HIF)-activated downstream gene and is normally responsible of the activation of mitochondrial proapoptotic cascades when overexpressed [70]. As energy production processes (protein and lipid oxidation, oxidative phosphorylation and related regulatory pathways) were enriched in embryos from nulliparous mares, it seemed unlikely that *FAM162A* up-regulation in embryos from nulliparous mares was a response to hypoxic environment. Nevertheless, it could be hypothesized that the uterine environment of embryos may vary according to mares’ parity, partly due to reduced uterine blood perfusion in nulliparous mares. To the authors’ knowledge, there is no study on the effect of nulliparity on uterine vascularization in young mares. It has nonetheless been shown that *FAM162A* expression is increased in intestinal and uterine cervical cancer [71, 72] its overexpression enhanced cell proliferation processes, suggesting a non-elucidated positive role in tumor development [72]. As in tumor, here, *FAM162A* could play a role in cell proliferation of equine embryos but the process remains to be elucidated. This could also indicate that proliferation in equine embryos differ according to mares’ parity, maybe as a response to their environment.

In the TE of embryos from nulliparous mares, pathways related to oxidative phosphorylation were enriched. The enrichment of the expression of genes involved in these pathways could indicate that the production of ATP from oxidative phosphorylation is up regulated in TE of embryos from nulliparous mares in comparison to the ones from multiparous mares. This up-regulation of oxygen oxidation in mitochondria could be harmful for TE cells as oxidative phosphorylation is accompanied by the production of reactive oxygen species (ROS) and particularly of hydrogen peroxide (for review [75]). Pathways related to hydrogen peroxide metabolic processes, however, were also enriched in the TE of embryos from nulliparous mares, showing that there is an up-regulation of the control of ROS such as hydrogen peroxide. The up regulation of both oxidative phosphorylation and regulation of ROS pathways suggests that there is an increased production of energy that is not harmful because well controlled in the TE of embryos from nulliparous mares.

At this developmental stage in equine embryos, 40 to 50% of glucose uptake is oxidized in the mitochondria, probably to meet the high energy demand of ionic transport associated with the important growth of both blastocoelic cavity and trophoblast [76]. Here, however, the enrichment in oxidative phosphorylation was not accompanied by an enrichment in glucose metabolism nor transport pathways but pathways linked to beta-oxidation of lipids and degradation of amino acids were enriched. This suggests that glycolysis is not affected by maternal parity but, to meet energy requirements, embryos from nulliparous mares use fatty acids and/or amino acids whilst embryos from multiparous mares do not need more energy than already provided and therefore, do not require the degradation of these substrates. The increased catabolism of amino acids and lipids could be detrimental for embryo development as the first are required for protein synthesis and the latter are mandatory for hormone production (for review [77, 78]). Pathways related to amino acid degradation, however, were mostly enriched in genes involved in beta-oxidation of fatty acids. Moreover, pathways related to translation and protein maturation but not pathways related to amino acids transport were enriched in embryos from nulliparous mares. Altogether, these results suggest that only lipid catabolism is enriched in the TE of nulliparous mares’ embryos.

In addition, pathways related to the transport of lipids and cholesterol were enriched in the TE of nulliparous mares’ embryos compared to those of multiparous mares. One hypothesis to explain these results could be that there is a higher energy demand in embryos from nulliparous mares and that they would compensate by degrading more lipids for oxidative phosphorylation, which requires more lipids to be obtained from the external environment. Another possibility is that the lipid composition of the uterine environment is altered in nulliparous mares, possibly due to immature uterine glands, leading to increased absorption by the embryo, that would stimulate beta-oxidation and thus oxidative phosphorylation. Indeed, the metabolism of lipids was also shown to be perturbed in blastocysts at the same developmental stage according to maternal parity in old mares [36]. To the authors’ knowledge, there is no study on the effects of maternal parity in any species on uterine fluid composition and how it could interfere with embryo gene expression. Although it is more likely that there are modifications in the uterine environment according to mare’s parity, the present results cannot conclude about the origin of the altered embryo metabolism.

As a confirmation of increased lipid transport, retinol binding protein 1 (*RBP1*) was up regulated in the ICM of embryos from nulliparous mares. Retinol is well known to be an important regulator of vertebrate development (for review [79]). In bovine, the addition of retinol to the maturation and culture medium of oocytes and embryos increased the blastocyst rate [80]. RBP transports the hydrophobic retinol in physiological fluids such as plasma [81] or uterine fluids. Pig conceptuses at the time of elongation produce RBP in large amounts, suggesting that retinol is important for embryo development [82]. In horses, the expression of *RBP* increased in the endometrium during diestrus under steroid regulation but did not vary according to the presence of an embryo or not [83]. Although underlying mechanisms are missing, *RBP1* could, however, play an important role in equine early embryo development by transporting retinol to the embryo. The increased expression of RBP in the ICM of embryos from nulliparous mares could be a response to a reduced availability of retinol in the close environment of the embryo or an increased requirements of retinol from embryos of nulliparous mares.

Nutrient and ion exchanges were also modified by maternal parity in the TE. Indeed, solute carrier family 47 member 1 (*SLC47A1* also known as *MATE1*), the solute transporter for molecules such as creatinine or guanidine, was down regulated in the TE of embryos from nulliparous mares. In addition, the expression of EF-hand calcium binding domain 11 (*EFCAB11*) and GTP binding protein 8 (*GTPBP8*) was increased in the TE of embryos from nulliparous mares compared to that of multiparous mares. These results could indicate a perturbed transport of different molecules in the TE of embryos from nulliparous mares.

These modifications of cell metabolism in the TE were associated with an alteration of pathways related to immunity, especially those linked to interleukin 1 beta (IL1B), being enriched in embryos from nulliparous mares. In cattle, it has been suggested that the early bovine embryo interacts with the dam’s immune system through processes involving IL1 [84]. In horses, maternal recognition of pregnancy (MRP) is thought to take place between 10-13 days post ovulation (for review [85]). At 19- and 25-days, but not at 13 days post ovulation, expression of the *IL1 receptor antagonist* has been shown to be markedly increased in the endometrium of pregnant compared to cyclic mares, suggesting that the endometrium regulates the IL1 signal and that IL1 plays a role in MRP in equine [86]. The expression of *IL1B* is increased in the luminal epithelium of pregnant *vs* cyclic mares at 10-13 days post ovulation, confirming the involvement of *IL1B* signaling process in MRP [87]. Here, embryos were collected earlier from the assumed MRP period but the observed differences in the IL1B signaling pathway could indicate that embryo-maternal communication and possibly MRP are affected by maternal parity.

Furthermore, related to lipid metabolism and IL1B signaling, peroxiredoxin like 2B (*PRXL2B*), also known as Prostamide/Prostaglandin F Synthase, was downregulated in the TE of nulliparous mares’ embryos. This gene encodes for an enzyme that has been shown to catalyze the reduction of prostamide H_2_ to prostamide F_2α_ as well as the reduction of prostaglandin H_2_ (PGH_2_) to prostaglandin F_2 α_ (PGF_2α_) [88]. In bovine, IL1B upregulates PGF_2α_ and prostamide secretion by in vitro cultured endometrial cells [89]. In horses, PGF_2α_ is secreted by the uterus to provoke the corpus luteum luteolysis (for review [90]). It has been shown that the suppression of the pulsatile secretion of PGF_2α_ from the endometrium is responsible for the maintenance of pregnancy [91] and that *in vitro*, PGF_2α_ production is significantly reduced when endometrial explants are co-cultured with embryonic tissues [92]. From the oviduct stages, equine embryos are able to produce prostaglandins [93–95]. Prostaglandins produced by the embryo, however, do not reach the blood circulation in sufficient amount to induce luteolysis [91]. It has been shown that these prostaglandins are required for myometrial contractions that participate in the migration of the equine embryo at the time of MRP [96]. By impeding the movement of the embryo, one study observed that equine embryo migration through at least 2/3 of the uterus is required to prevent luteolysis [97]. Moreover, the use of an intra-uterine device to imitate the physical presence of an embryo, allowed to prevent the luteolysis [98]. A recent study, nevertheless, observed that the contact of a substance/object is not sufficient to reduce PGF secretion from the endometrium, suggesting that embryo secretions are required for luteolysis [99]. Therefore, although MRP is thought to begin 2 days later, the present study shows that MRP might be delayed or disturbed in nulliparous mares.

## 5. Conclusion

So far, the effect of mare’s parity on embryo gene expression had never been considered. The present study shows that mare’s parity affects the expression of genes in both ICM and TE of blastocysts. Only the expression of few genes is altered but several important functions for embryo development are affected by mare’s parity. Indeed, nulliparity in young mares particularly alters the expression of genes related to transcription and RNA processing in the ICM and embryo-maternal communication in the TE, suggesting embryo adaptation to an environment that is different in nulliparous *vs* multiparous mares. Individual chances of implantation for each embryo could not be predicted by the results of this study. Until today, the capacity of uterus to enlarge and support pregnancy was the only suggested explanation for the lighter and smaller foal and placenta at birth in nulliparous mares. The present results indicate differences in embryo-maternal communication long before implantation that could alter the embryo development as well as maternal recognition of pregnancy.

## Data Availability Statement

The RNA sequençing data supporting the conclusions of this article are available in the GEO SuperSeries [accession: GSE193676; https://www.ncbi.nlm.nih.gov/geo/query/acc.cgi?acc=GSE193676], containing repositories [accession: GSE162893; https://www.ncbi.nlm.nih.gov/geo/query/acc.cgi?acc=GSE162893] and [accession: GSE193675; https://www.ncbi.nlm.nih.gov/geo/query/acc.cgi?acc=GSE193675].

## Acknowledgments

The authors are grateful to the staff of the Institut Français du Cheval et de l’Equitation (IFCE) experimental farm (Plateau technique de la Valade, Chamberet, France) for care and management of animals. We acknowledge the high-throughput sequencing facility of I2BC for its sequencing and bioinformatics expertise. The bioinformatics analyses were performed thanks to Core Cluster of the Institut Français de Bioinformatique (IFB) (ANR-11-INBS-0013). Many thanks to Matthias Zytnicki and Christophe Klopp for their advice on RNA-seq de novo analysis. Many thanks to Pablo Ross who kindly provided the coordinates for the XIST gene.

## Author contributions

PCP obtained the funding. PCP and VD conceived the project. VD and PCP supervised the study. ED, CA, ND, NP, VD and PCP adapted the methodology for the project. ED, CA, JAR and YJ performed the experiments. CA, ND, NP and MD provided the resources. ED, LJ, YJ and RL performed data curation. ED and LJ analyzed the data. ED wrote the original draft. All authors read, revised, and approved the submitted manuscript.

## List of abbreviations

DEG: differential expressed genes
DeMixT_ICM_cells: deconvoluted gene expression in ICM cells
DeMixT_TE_cells: deconvoluted gene expression in TE cells
ECM: Extracellular matrix
ERR: embryo collection rate
FDR: false discovery rate
GO BP: Gene Ontology biological process
GO: Gene Ontology
GSEA: gene set enrichment analyses
ICM: inner cell mass
ICMandTE: inner cell mass enriched hemi-embryo
ICSI: intracytoplasmic sperm injection
IL1B: Interleukin 1 beta
KEGG: Kyoto Encyclopedia of Genes and Genomes
Log2FC: log2 fold change
NES: normalized enrichment score
OM: old multiparous mares
ON: old nulliparous mares
TE: trophoblast
TE_part: pure trophoblast hemi-embryo
XIST: X inactive Specific Transcript

## Supplementary material

Supplementary Figure 1:

SupFig1_Embryosize.tif

Plot of equine individual embryo according to their size

Supplementary Figure 2:

SupFig2_comp_with_without.png

Venn diagrams of differential analyses on equine embryo gene expression according to maternal parity in the inner cell mass (ICM) and the trophoblast part (TE) with or without the largest embryo (2643µm in diameter)

Supplementary Table 1:

Sup1_ICM_Diff_avecYME5.csv

Differential gene analysis using DeSeq2 in DeMixT_ICM_cells of equine embryo at Day 8 post-ovulation according to mares’ parity with the large embryo

Equine ensemble ID, orthologue human Ensembl ID, Orthologue human Entrez Gene ID, gene description, normalized counts for each embryo and parameters obtained after Deseq2 analysis (log2FoldChange, pvalue and padj (after FDR correction)) of genes expressed in ICM (after gene expression deconvolution of ICMandTE using DeMixT) of YN and YM embryos

ICM: Inner cell mass; YN: young nulliparous mares; YM: young multiparous mares

Supplementary Table 2:

Sup2_TE_Diff_avecYME5.csv

Differential gene analysis using DeSeq2 in TE_part of equine embryo at Day 8 post-ovulation according to mares’ parity with the large embryo

Equine ensemble ID, orthologue human Ensembl ID, Orthologue human Entrez Gene ID, gene description, normalized counts for each embryo and parameters obtained after Deseq2 analysis (log2FoldChange, pvalue and padj (after FDR correction)) of genes expressed in TE_part of YN and YM embryos

TE: trophoblast; YN: young nulliparous mares; YM: young multiparous mares

Supplementary Table 3:

Sup3_ICM_Diff_sansYME5.csv

Differential gene analysis using DeSeq2 in DeMixT_ICM_cells of equine embryo at Day 8 post-ovulation according to mares’ parity without the largest embryo

Equine ensemble ID, orthologue human Ensembl ID, Orthologue human Entrez Gene ID, gene description, normalized counts for each embryo and parameters obtained after Deseq2 analysis (log2FoldChange, pvalue and padj (after FDR correction)) of genes expressed in ICM (after gene expression deconvolution of ICMandTE using DeMixT) of YN and YM embryos

ICM: Inner cell mass; YN: young nulliparous mares; YM: young multiparous mares

Supplementary Table 4:

Sup4_TE_Diff_sansYME5.csv

Differential gene analysis using DeSeq2 in TE_part of equine embryo at Day 8 post-ovulation according to mares’ parity with the largest embryo

Equine ensemble ID, orthologue human Ensembl ID, Orthologue human Entrez Gene ID, gene description, normalized counts for each embryo and parameters obtained after Deseq2 analysis (log2FoldChange, pvalue and padj (after FDR correction)) of genes expressed in TE_part of YN and YM embryos

TE: trophoblast; YN: young nulliparous mares; YM: young multiparous mares

Supplementary Table 5:

Sup5_ICM_GSEA_sansYME5.csv

Gene set enrichment analysis results on gene expression of DeMixT_ICM_cells of embryos from young nulliparous and multiparous mares

Gene Set Enrichment Analysis results (database, pathway name, size, enrichment score without and with normalization, p-value and FDR corrected q-value) for GO biological process, KEGG and REACTOME databases on DeMixT_ICM_cells gene expression table. These results did not include YM_E5, the embryo larger than 2,000µm. ICM: Inner cell mass

Supplementary Table 6:

Sup6_TE_GSEA_sansYME5.csv

Gene set enrichment analysis results on gene expression of TE_part of embryos from young nulliparous and multiparous mares

Gene Set Enrichment Analysis results (database, pathway name, size, enrichment score without and with normalization, p-value and FDR corrected q-value) for GO biological process, KEGG and REACTOME databases on TE_part gene expression table. These results did not include YM_E5, the embryo larger than 2,000µm.

## References

[1] Roseboom T, de Rooij S, Painter R. The Dutch famine and its long-term consequences for adult health. Early Human Development 2006;82:485–91. https://doi.org/10.1016/j.earlhumdev.2006.07.001.

[2] Chavatte-Palmer P, Velazquez MA, Jammes H, Duranthon V. Review: Epigenetics, developmental programming and nutrition in herbivores 2018:1–9. https://doi.org/10.1017/S1751731118001337.

[3] Doreau M, Boulot S, Martin-Rosset W. Effect of parity and physiological state on intake, milk production and blood parameters in lactating mares differing in body size. Animal Production 1991;53:111–8. https://doi.org/10.1017/S0003356100006048.

[4] Lawrence LM, DiPietro J, Ewert K, Parrett D, Moser L, Powell D. Changes in body weight and condition of gestating mares. Journal of Equine Veterinary Science, 1992;12:355--358. https://doi.org/10.1016/S0737-0806(06)81361-4.

[5] Pool-Anderson K, Raub RH, Warren JA. Maternal Influences on Growth and Development of Full-Sibling Foals. Journal of Animal Science 1994;72:1661–6.

[6] Cymbaluk NF, Laarveld B. The ontogeny of serum insulin-like growth factor-I concentration in foals: effects of dam parity, diet, and age at weaning. Domestic Animal Endocrinology 1996;13:197–209. https://doi.org/0739724096000148 [pii].

[7] Wilsher S, Allen WR. The effects of maternal age and parity on placental and fetal development in the mare. Equine Veterinary Journal 2003;35:476–83. https://doi.org/10.2746/042516403775600550.

[8] Elliott C, Morton J, Chopin J. Factors affecting foal birth weight in Thoroughbred horses. Theriogenology 2009;71:683–9. https://doi.org/10.1016/j.theriogenology.2008.09.041.

[9] Fernandes CB, Meirelles MG, Guimaraes CF, Nichi M, Affonso FJ, Fonte JS, et al. Which paternal, maternal and placental parameters influence foal size and vitality? Journal of Equine Veterinary Science 2014;34:225–7. https://doi.org/10.1016/j.jevs.2013.10.161.

[10] Klewitz J, Struebing C, Rohn K, Goergens A, Martinsson G, Orgies F, et al. Effects of age, parity, and pregnancy abnormalities on foal birth weight and uterine blood flow in the mare. Theriogenology 2015;83:721–9. https://doi.org/10.1016/j.theriogenology.2014.11.007.

[11] Affonso FJ, Meirelles MG, Alonso MA, Guimaraes CF, Lemes KM, Nichi M, et al. Influence of mare parity on weight, height, thoracic circumference and vitality of neonatal foals. Journal of Equine Veterinary Science 2016;41:67. https://doi.org/10.1016/j.jevs.2016.04.053.

[12] Meirelles MG, Veras MM, Alonso MA, de Fátima Guimarães C, Nichi M, Fernandes CB. Influence of maternal age and parity on placental structure and foal characteristics from birth up to two years of age. Journal of Equine Veterinary Science 2017. https://doi.org/10.1016/j.jevs.2017.03.226.

[13] Robles M, Dubois C, Gautier C, Dahirel M, Guenon I, Bouraima-Lelong H, et al. Maternal parity affects placental development, growth and metabolism of foals until 1 year and a half. Theriogenology 2018;108:321–30. https://doi.org/10.1016/j.theriogenology.2017.12.019.

[14] Barron JK. The effect of maternal age and parity on the racing performance of Thoroughbred horses. Equine Veterinary Journal 1995;27:73–5. https://doi.org/10.1111/j.2042-3306.1995.tb03036.x.

[15] Palmer E, Robles M, Chavatte-Palmer P, Ricard A. Maternal Effects on Offspring Performance in Show Jumping. Journal of Equine Veterinary Science 2018;66:221. https://doi.org/10.1016/j.jevs.2018.05.108.

[16] Wilsher S, Allen WR. The effects of maternal age and parity on placental and fetal development in the mare. Equine Veterinary Journal 2003;35:476–83. https://doi.org/10.2746/042516403775600550.

[17] Allen WR, Stewart F. Equine placentation. Reprod Fertil Dev 2001;13:623. https://doi.org/10.1071/RD01063.

[18] Steven DH, Samuel CA. Anatomy of the placental barrier in the mare. Journal of Reproduction and Fertility Supplement 1975;23:579–82.

[19] Samuel CA, Allen WR, Steven DH. Studies on the equine placenta II. Ultrastructure of the placental barrier. Journal of Reproduction and Fertility Supplement 1976;48:257–64.

[20] Allen WR, Wilsher S. A Review of Implantation and Early Placentation in the Mare. Placenta 2009;30:1005–15. https://doi.org/10.1016/j.placenta.2009.09.007.

[21] Platt H. Aetiological aspects of abortion in the thoroughbred mare. J Comp Pathol 1973;83:199– 205.

[22] Chevalier-Clément F. Pregnancy loss in the mare. Animal Reproduction Science 1989;20:231– 44. https://doi.org/10.1016/0378-4320(89)90088-2.

[23] Gibbs PG, Davison KE. A field study on reproductive efficiency of mares maintained predominately on native pasture. Journal of Equine Veterinary Science 1992;12:219–22. https://doi.org/10.1016/S0737-0806(06)81449-8.

[24] Vidament M, Dupere AM, Julienne P, Evain A, Noue P, Palmer E. Equine frozen semen: freezability and fertility field results. THERIOGENOLOGY 1997;48:907–17. https://doi.org/10.1016/S0093-691X(97)00319-1.

[25] Morris LHA, Allen WR. Reproductive efficiency of intensively managed Thoroughbred mares in Newmarket. Equine Veterinary Journal 2002;34:51–60. https://doi.org/10.2746/042516402776181222.

[26] Langlois B, Blouin C. Statistical analysis of some factors affecting the number of horse births in France. Reprod Nutr Dev 2004;44:583–95. https://doi.org/10.1051/rnd:2004055.

[27] Hanlon D, Stevenson M, Evans M, Firth E. Reproductive performance of Thoroughbred mares in the Waikato region of New Zealand: 1. Descriptive analyses. New Zealand Veterinary Journal 2012;60:329–34. https://doi.org/10.1080/00480169.2012.693039.

[28] Baker CB, Little TV, McDOWELL KJ. The live foaling rate per cycle in mares. Equine Veterinary Journal 1993;25:28–30. https://doi.org/10.1111/j.2042-3306.1993.tb04819.x.

[29] Barbacini S, Marchi V, Zavaglia G. Equine frozen semen: results obtained in Italy during the 1994-1997 period. Equine Veterinary Education 1999;11:109–12. https://doi.org/10.1111/j.2042-3292.1999.tb00930.x.

[30] Brück I, Anderson G, Hyland J. Reproductive performance of Thoroughbred mares on six commercial stud farms. Australian Vet J 1993;70:299–303. https://doi.org/10.1111/j.1751-0813.1993.tb07979.x.

[31] Hemberg E, Lundeheim N, Einarsson S. Reproductive Performance of Thoroughbred Mares in Sweden. Reprod Domest Anim 2004;39:81–5. https://doi.org/10.1111/j.1439-0531.2004.00482.x.

[32] Nath L, Anderson G, McKinnon A. Reproductive efficiency of Thoroughbred and Standardbred horses in north-east Victoria. Australian Veterinary Journal 2010;88:169–75. https://doi.org/10.1111/j.1751-0813.2010.00565.x.

[33] Carluccio A, Bucci R, Fusi J, Robbe D, Veronesi MC. Effect of age and of reproductive status on reproductive indices in horse mares carrying mule pregnancies. Heliyon 2020;6:e05175. https://doi.org/10.1016/j.heliyon.2020.e05175.

[34] Derisoud E, Auclair-Ronzaud J, Palmer E, Robles M, Chavatte-Palmer P, Derisoud E, et al. Female age and parity in horses: how and why does it matter? Reprod Fertil Dev 2021;34:52–116. https://doi.org/10.1071/RD21267.

[35] Derisoud E, Jouneau L, Dubois C, Archilla C, Jaszczyszyn Y, Legendre R, et al. Maternal age affects equine Day 8 embryo gene expression both in trophoblast and inner cell mass. BioRxiv 2021:2021.04.07.438786. https://doi.org/10.1101/2021.04.07.438786.

[36] Derisoud E, Jouneau L, Gourtay C, Margat A, Archilla C, Jaszczyszyn Y, et al. Maternal parity affects Day 8 embryo gene expression in old mares. BioRxiv 2021. https://doi.org/10.1101/2021.12.01.470709.

[37] Duchamp G, Bour B, Combarnous Y, Palmer E. Alternative solutions to hCG induction of ovulation in the mare. J Reprod Fertil Suppl 1987;35:221–8.

[38] Bucca S, Carli A. Efficacy of human chorionic gonadotropin to induce ovulation in the mare, when associated with a single dose of dexamethasone administered at breeding time: Efficacy of human chorionic gonadotropin to induce ovulation when associated with dexamethasone. Equine Veterinary Journal 2011;43:32–4. https://doi.org/10.1111/j.2042-3306.2011.00488.x.

[39] Enders AC, Schlafke S, Lantz KC, Liu IKM. Endoderm cells of the equine yolk sac from Day 7 until formation of the definitive yolk sac placenta. Equine Veterinary Journal 1993;25:3–9. https://doi.org/10.1111/j.2042-3306.1993.tb04814.x.

[40] Enders AC, Lantz KC, Liu IKM, Schlafke S. Loss of polar trophoblast during differentiation of the blastocyst of the horse. Journal of Reproduction and Fertility 1988;83:447–60. https://doi.org/10.1530/jrf.0.0830447.

[41] Martin M. Cutadapt removes adapter sequences from high-throughput sequencing reads. EMBnet Journal 2011;17:10–2.

[42] Dobin A, Davis CA, Schlesinger F, Drenkow J, Zaleski C, Jha S, et al. STAR: ultrafast universal RNA-seq aligner. Bioinformatics 2013;29:15–21. https://doi.org/10.1093/bioinformatics/bts635.

[43] Liao Y, Smyth GK, Shi W. featureCounts: an efficient general purpose program for assigning sequence reads to genomic features. Bioinformatics 2014;30:923–30. https://doi.org/10.1093/bioinformatics/btt656.

[44] R Core Team. A Language and Environment for Statistical Computing. Vienna, Austria: R foundation for Statistical Computing; 2020.

[45] Rstudio Team. RStudio: Integrated Development for R. RStudio. Boston, USA: PBC; 2020.

[46] Pinheiro J, Bates D, Debroy S, Sarkar D, R Core Team. nlme: Linear and Nonlinear Mixed Effects Models. 2020.

[47] Giraudoux P. pgirmess: Spatial Analysis and Data Mining for Field Ecologists. 2018.

[48] Cao S, Wang JR, Ji S, Yang P, Chen J, Shen JP, et al. Differing total mRNA expression shapes the molecular and clinical phenotype of cancer. BioRxiv 2020:57. https://doi.org/10.1101/2020.09.30.306795.

[49] Wang Z, Cao S, Morris JS, Ahn J, Liu R, Tyekucheva S, et al. Transcriptome Deconvolution of Heterogeneous Tumor Samples with Immune Infiltration. IScience 2018;9:451–60. https://doi.org/10.1016/j.isci.2018.10.028.

[50] Love MI, Huber W, Anders S. Moderated estimation of fold change and dispersion for RNA-seq data with DESeq2. Genome Biol 2014;15:550. https://doi.org/10.1186/s13059-014-0550-8.

[51] Iqbal K, Chitwood JL, Meyers-Brown GA, Roser JF, Ross PJ. RNA-Seq Transcriptome Profiling of Equine Inner Cell Mass and Trophectoderm. Biology of Reproduction 2014;90. https://doi.org/10.1095/biolreprod.113.113928.

[52] Gómez-Rubio V. ggplot2 - Elegant Graphics for Data Analysis (2nd Edition). J Stat Soft 2017;77. https://doi.org/10.18637/jss.v077.b02.

[53] Kassambara A, Mundt F. factoextra: Extract and Visualize the Results of Multivariate Data Analyses. 2020.

[54] Kolberg L, Raudvere U. gprofiler2: Interface to the “g:Profiler” Toolset. 2020.

[55] Yates AD, Achutan P, Akanni W, Allen J, Allen J, Alvarez-Jarreta J, et al. Ensembl 2020 n.d.

[56] Subramanian A, Tamayo P, Mootha VK, Mukherjee S, Ebert BL, Gillette MA, et al. Gene set enrichment analysis: A knowledge-based approach for interpreting genome-wide expression profiles. Proceedings of the National Academy of Sciences 2005;102:15545–50. https://doi.org/10.1073/pnas.0506580102.

[57] Mootha VK, Lindgren CM, Eriksson K-F, Subramanian A, Sihag S, Lehar J, et al. PGC-1α-responsive genes involved in oxidative phosphorylation are coordinately downregulated in human diabetes. Nat Genet 2003;34:267–73. https://doi.org/10.1038/ng1180.

[58] Liberzon A, Birger C, Thorvaldsdóttir H, Ghandi M, Mesirov JP, Tamayo P. The Molecular Signatures Database Hallmark Gene Set Collection. Cell Systems 2015;1:417–25. https://doi.org/10.1016/j.cels.2015.12.004.

[59] Savage SR, Shi Z, Liao Y, Zhang B. Graph Algorithms for Condensing and Consolidating Gene Set Analysis Results. Molecular & Cellular Proteomics 2019;18:S141–52. https://doi.org/10.1074/mcp.TIR118.001263.

[60] Shannon P. Cytoscape: A Software Environment for Integrated Models of Biomolecular Interaction Networks. Genome Research 2003;13:2498–504. https://doi.org/10.1101/gr.1239303.

[61] McKinnon AO, Squires EL. Morphologic assessment of the equine embryo. Journal of the American Veterinary Medical Association 1988;192:401–6.

[62] Derisoud E, Jouneau L, Margat A, Gourtay C, Dubois C, Archilla C, et al. 52 Equine embryo size does matter! Reprod Fertil Dev 2021;34:261–261. https://doi.org/10.1071/RDv34n2Ab52.

[63] Tucker RP, Kenzelmann D, Trzebiatowska A, Chiquet-Ehrismann R. Teneurins: Transmembrane proteins with fundamental roles in development. The International Journal of Biochemistry & Cell Biology 2007;39:292–7. https://doi.org/10.1016/j.biocel.2006.09.012.

[64] Vanderwall DK. Early Embryonic Development and Evaluation of Equine Embryo Viability. Veterinary Clinics of North America: Equine Practice 1996;12:61–83. https://doi.org/10.1016/S0749-0739(17)30295-X.

[65] Panzani D, Rota A, Marmorini P, Vannozzi I, Camillo F. Retrospective study of factors affecting multiple ovulations, embryo recovery, quality, and diameter in a commercial equine embryo transfer program. Theriogenology 2014;82:807–14. https://doi.org/10.1016/j.theriogenology.2014.06.020.

[66] Betteridge KJ. The structure and function of the equine capsule in relation to embryo manipulation and transfer. Equine Veterinary Journal 2010;21:92–100. https://doi.org/10.1111/j.2042-3306.1989.tb04690.x.

[67] Beckelmann J, Budik S, Bartel C, Aurich C. Evaluation of Xist expression in preattachment equine embryos. Theriogenology 2012;78:1429–36. https://doi.org/10.1016/j.theriogenology.2012.05.026.

[68] Colchen S, Battut I, Fiéni F, Tainturier D, Siliart B, Bruyas JF. Quantitative histological analysis of equine embryos at exactly 156 and 168 h after ovulation. J Reprod Fertil Suppl 2000:527–37.

[69] Leisinger CA, Medina V, Markle ML, Paccamonti DL, Pinto CRF. Morphological evaluation of Day 8 embryos developed during induced aluteal cycles in the mare. Theriogenology 2018;105:178– 83. https://doi.org/10.1016/j.theriogenology.2017.09.029.

[70] Lee M-J, Kim J-Y, Suk K, Park J-H. Identification of the Hypoxia-Inducible Factor 1α-Responsive HGTD-P Gene as a Mediator in the Mitochondrial Apoptotic Pathway. Mol Cell Biol 2004;24:3918–27. https://doi.org/10.1128/MCB.24.9.3918-3927.2004.

[71] Cho Y-E, Kim J-Y, Kim Y-J, Kim Y-W, Lee S, Park J-H. Expression and clinicopathological significance of human growth and transformation-dependent protein (HGTD-P) in uterine cervical cancer. Histopathology 2010;57:479–82. https://doi.org/10.1111/j.1365-2559.2010.03627.x.

[72] Cho Y-E, Kim J-Y, Kim Y-W, Park J-H, Lee S. Expression and prognostic significance of human growth and transformation-dependent protein in gastric carcinoma and gastric adenoma. Human Pathology 2009;40:975–81. https://doi.org/10.1016/j.humpath.2008.12.007.

[73] Charpentier AH, Bednarek AK, Daniel RL, Hawkins KA, Laflin KJ, Gaddis S, et al. Effects of Estrogen on Global Gene Expression: Identification of Novel Targets of Estrogen Action. Cancer Res 2000;60:5977–83.

[74] Ao A, Wang H, Kamarajugadda S, Lu J. Involvement of estrogen-related receptors in transcriptional response to hypoxia and growth of solid tumors. PNAS 2008;105:7821–6. https://doi.org/10.1073/pnas.0711677105.

[75] Balaban RS, Nemoto S, Finkel T. Mitochondria, Oxidants, and Aging. Cell 2005;120:483–95. https://doi.org/10.1016/j.cell.2005.02.001.

[76] Lane M, O’Donovan MK, Squires EL, Seidel GE, Gardner DK. Assessment of metabolism of equine morulae and blastocysts. Mol Reprod Dev 2001;59:33–7. https://doi.org/10.1002/mrd.1004.

[77] Leese HJ, McKeegan PJ, Sturmey RG. Amino Acids and the Early Mammalian Embryo: Origin, Fate, Function and Life-Long Legacy. International Journal of Environmental Research and Public Health 2021;18:9874. https://doi.org/10.3390/ijerph18189874.

[78] Sturmey R, Reis A, Leese H, McEvoy T. Role of Fatty Acids in Energy Provision During Oocyte Maturation and Early Embryo Development. Reproduction in Domestic Animals 2009;44:50–8. https://doi.org/10.1111/j.1439-0531.2009.01402.x.

[79] Marceau G, Gallot D, Lemery D, Sapin V. Metabolism of Retinol During Mammalian Placental and Embryonic Development. Vitamins & Hormones, vol. 75, Academic Press; 2007, p. 97–115. https://doi.org/10.1016/S0083-6729(06)75004-X.

[80] Livingston T, Eberhardt D, Edwards JL, Godkin J. Retinol improves bovine embryonic development in vitro. Reprod Biol Endocrinol 2004;2:83. https://doi.org/10.1186/1477-7827-2-83.

[81] Goodman DS. 8 - Plasma Retinol-Binding Protein. In: Sporn MB, Roberts AB, Goodman DS, editors. The Retinoids, Academic Press; 1984, p. 41–88. https://doi.org/10.1016/B978-0-12-658102-7.50008-7.

[82] Trout WE, McDonnell JJ, Kramer KK, Baumbach GA, Roberts RM. The Retinol-Binding Protein of the Expanding Pig Blastocyst: Molecular Cloning and Expression in Trophectoderm and Embryonic Disc. Molecular Endocrinology 1991;5:1533–40. https://doi.org/10.1210/mend-5-10-1533.

[83] McDowell KJ, Adams MH, Franklin KM, Baker CB. Changes in Equine Endometrial Retinol-Binding Protein RNA during the Estrous Cycle and Early Pregnancy and with Exogenous Steroids1. Biology of Reproduction 1995;52:438–43. https://doi.org/10.1095/biolreprod52.2.438.

[84] Correia-Álvarez E, Gómez E, Martín D, Carrocera S, Pérez S, Otero J, et al. Expression and localization of interleukin 1 beta and interleukin 1 receptor (type I) in the bovine endometrium and embryo. Journal of Reproductive Immunology 2015;110:1–13. https://doi.org/10.1016/j.jri.2015.03.006.

[85] Swegen A. Maternal recognition of pregnancy in the mare: does it exist and why do we care? Reproduction 2021;161:R139–55. https://doi.org/10.1530/REP-20-0437.

[86] Haneda S, Nagaoka K, Nambo Y, Kikuchi M, Nakano Y, Matsui M, et al. Interleukin-1 receptor antagonist expression in the equine endometrium during the peri-implantation period. Domestic Animal Endocrinology 2009;36:209–18. https://doi.org/10.1016/j.domaniend.2008.11.006.

[87] Rudolf Vegas A, Podico G, Canisso IF, Bollwein H, Almiñana C, Bauersachs S. Spatiotemporal endometrial transcriptome analysis revealed the luminal epithelium as key player during initial maternal recognition of pregnancy in the mare. Sci Rep 2021;11:22293. https://doi.org/10.1038/s41598-021-01785-3.

[88] Moriuchi H, Koda N, Okuda-Ashitaka E, Daiyasu H, Ogasawara K, Toh H, et al. Molecular Characterization of a Novel Type of Prostamide/Prostaglandin F Synthase, Belonging to the Thioredoxin-like Superfamily *. Journal of Biological Chemistry 2008;283:792–801. https://doi.org/10.1074/jbc.M705638200.

[89] Almughlliq FB, Koh YQ, Peiris HN, Vaswani K, Arachchige BJ, Reed S, et al. Eicosanoid pathway expression in bovine endometrial epithelial and stromal cells in response to lipopolysaccharide, interleukin 1 beta, and tumor necrosis factor alpha. Reproductive Biology 2018;18:390–6. https://doi.org/10.1016/j.repbio.2018.10.001.

[90] Daels PF, Stabenfeldt GH, Kindahl H, Hughes JP. Prostaglandin release and luteolysis associated with physiological and pathological conditions of the reproductive cycle of the mare: a review. Equine Veterinary Journal 1989;21:29–34. https://doi.org/10.1111/j.2042-3306.1989.tb04669.x.

[91] Kindahl H, Knudsen O, Madej A, Edqvist LE. Progesterone, prostaglandin F-2 alpha, PMSG and oestrone sulphate during early pregnancy in the mare. J Reprod Fertil Suppl 1982;32:353–9.

[92] Berglund LA, Sharp DC, Vernon MW, Thatcher WW. Effect of pregnancy and collection technique on prostaglandin F in the uterine lumen of Pony mares. J Reprod Fertil Suppl 1982;32:335–41.

[93] Watson ED, Sertich PL. Prostaglandin production by horse embryos and the effect of co-culture of embryos with endometrium from pregnant mares. Reproduction 1989;87:331–6. https://doi.org/10.1530/jrf.0.0870331.

[94] Weber JA, Freeman DA, Vanderwall DK, Woods GL. Prostaglandin E2 secretion by oviductal transport-stage equine embryos. Biology of Reproduction 1991;45:540–3.

[95] Vanderwall DK, Woods GL, Weber JA, Lichtenwalner AB. Uterine transport of prostaglandin E2-releasing simulated embryonic vesicles in mares. Theriogenology 1993;40:13–20. https://doi.org/10.1016/0093-691X(93)90337-5.

[96] Stout T, Allen W. Role of prostaglandins in intrauterine migration of the equine conceptus. Reproduction 2001:771–5. https://doi.org/10.1530/rep.0.1210771.

[97] McDowell KJ, Sharp DC, Grubaugh W, Thatcher WW, Wilcox CJ. Restricted conceptus mobility results in failure of pregnancy maintenance in mares. Biology of Reproduction 1988;39:340–8.

[98] Rivera del Alamo MM, Reilas T, Kindahl H, Katila T. Mechanisms behind intrauterine device-induced luteal persistence in mares. Animal Reproduction Science 2008;107:94–106. https://doi.org/10.1016/j.anireprosci.2007.06.010.

[99] Klohonatz KM, Nulton LC, Hess AM, Bouma GJ, Bruemmer JE. The role of embryo contact and focal adhesions during maternal recognition of pregnancy. PLOS ONE 2019;14:e0213322. https://doi.org/10.1371/journal.pone.0213322.

