## Supplementary figures and images for "Nulliparity affects the expression of a limited number of genes and pathways in Day 8 equine embryos"

### Supplementary Figure 1

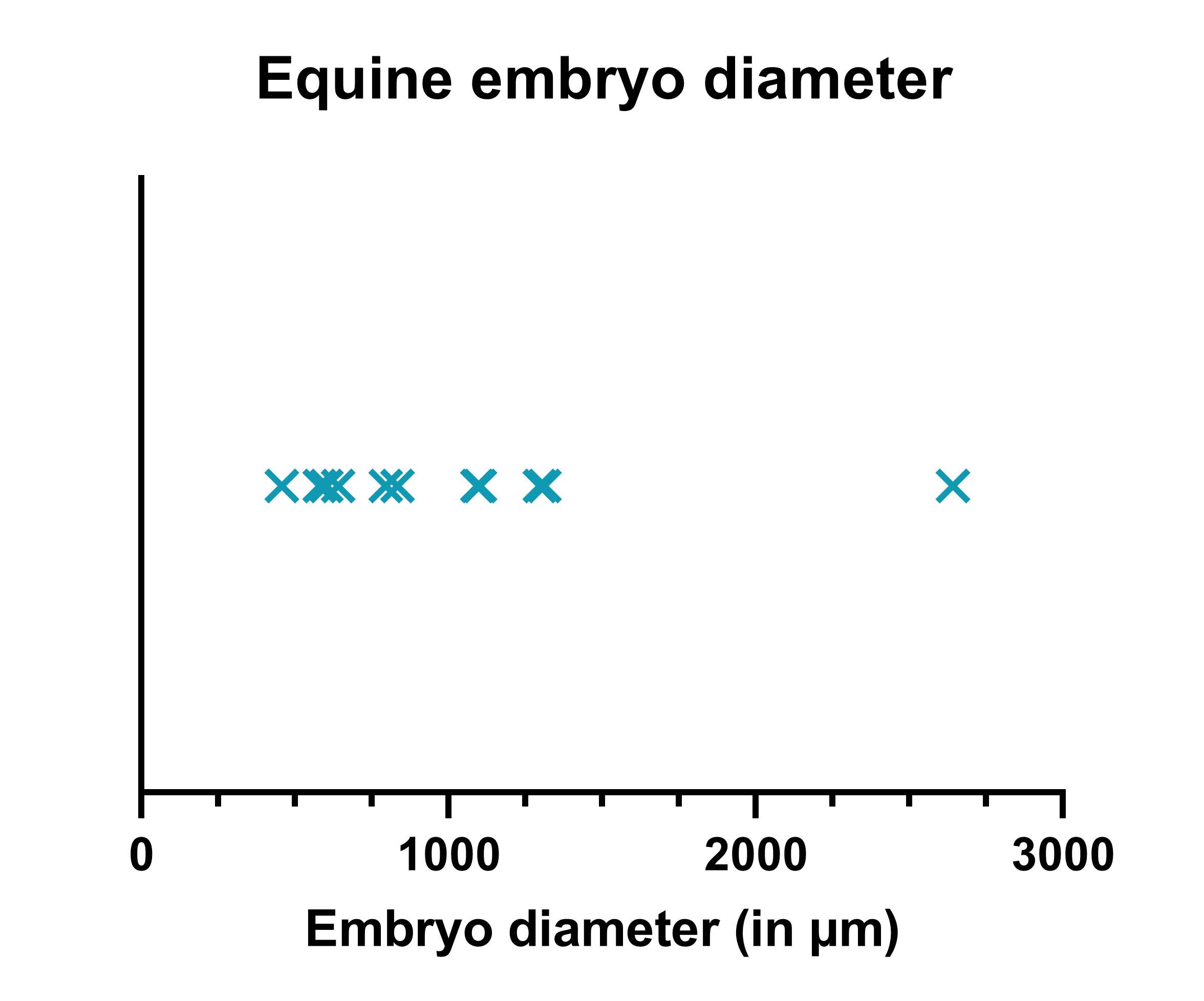
