## Supplementary Figure 2 for "Nulliparity affects the expression of a limited number of genes and pathways in Day 8 equine embryos"

### Slide 1
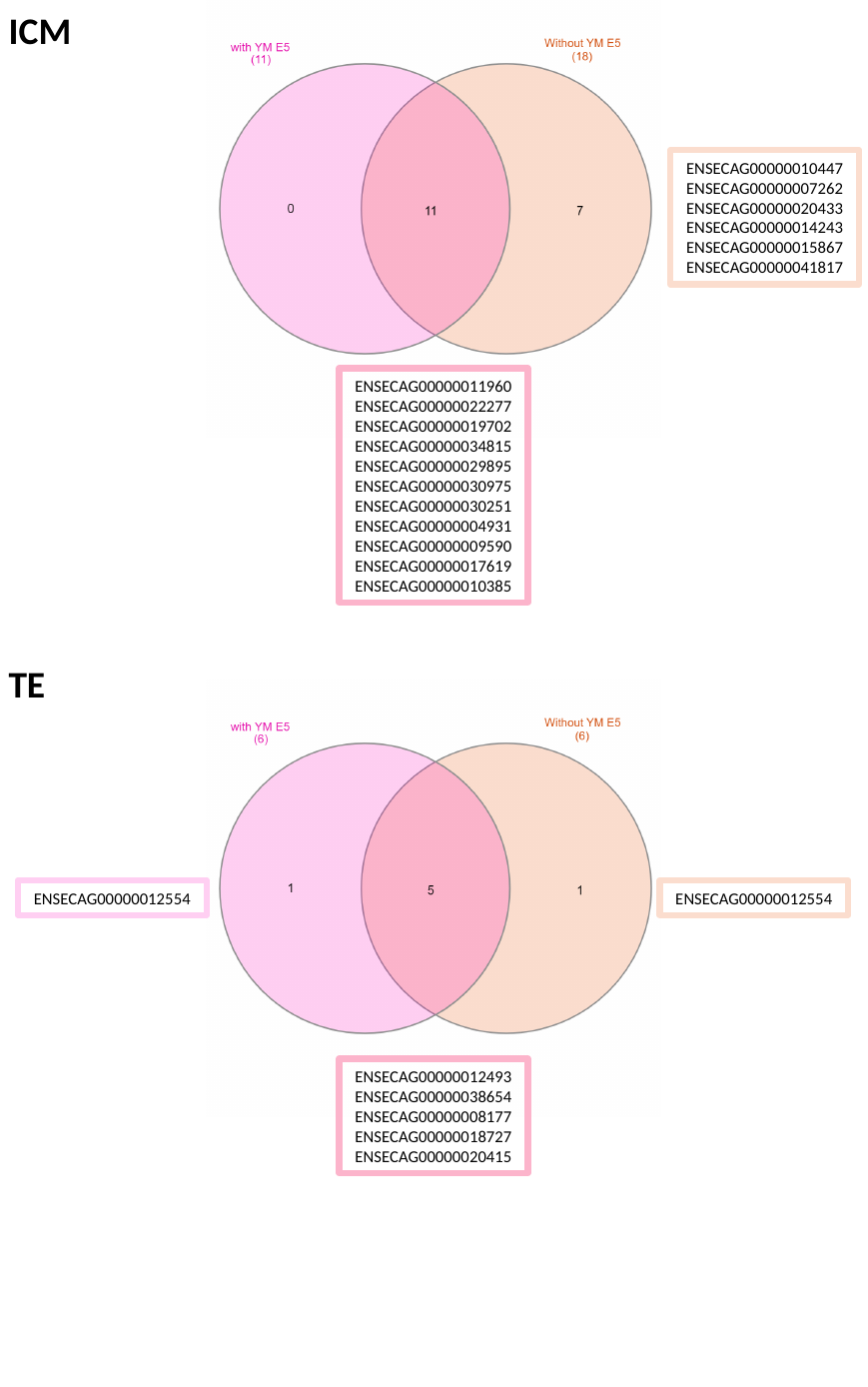

ICM
ENSECAG00000010447
ENSECAG00000007262
ENSECAG00000020433
ENSECAG00000014243
ENSECAG00000015867
ENSECAG00000041817
ENSECAG00000011960
ENSECAG00000022277
ENSECAG00000019702
ENSECAG00000034815
ENSECAG00000029895
ENSECAG00000030975
ENSECAG00000030251
ENSECAG00000004931
ENSECAG00000009590
ENSECAG00000017619
ENSECAG00000010385
TE
ENSECAG00000012554
ENSECAG00000012554
ENSECAG00000012493
ENSECAG00000038654
ENSECAG00000008177
ENSECAG00000018727
ENSECAG00000020415
